# Msh3 and Pms1 Set Neuronal CAG-repeat Migration Rate to Drive Selective Striatal and Cortical Pathogenesis in HD Mice

**DOI:** 10.1101/2024.07.09.602815

**Authors:** Nan Wang, Shasha Zhang, Peter Langfelder, Lalini Ramanathan, Mary Plascencia, Fuying Gao, Raymond Vaca, Xiaofeng Gu, Linna Deng, Leonardo E. Dionisio, Brinda C. Prasad, Thomas Vogt, Steve Horvath, Jeffrey S. Aaronson, Jim Rosinski, X. William Yang

## Abstract

Modifiers of Huntington’s disease (HD) include mismatch repair (MMR) genes; however, their underlying disease-altering mechanisms remain unresolved. Knockout (KO) alleles for 9 HD GWAS modifiers/MMR genes were crossed to the Q140 Huntingtin (mHtt) knock-in mice to probe such mechanisms. Four KO mice strongly (*Msh3* and *Pms1*) or moderately (*Msh2* and *Mlh1*) rescue a triad of adult-onset, striatal medium-spiny-neuron (MSN)-selective phenotypes: somatic *Htt* DNA CAG-repeat expansion, transcriptionopathy, and mHtt protein aggregation. Comparatively, Q140 cortex also exhibits an analogous, but later-onset, pathogenic triad that is *Msh3*-dependent. Remarkably, Q140/homozygous Msh3-KO lacks visible mHtt aggregates in the brain, even at advanced ages (20-months). Moreover, *Msh3*-deficiency prevents striatal synaptic marker loss, astrogliosis, and locomotor impairment in HD mice. Purified Q140 MSN nuclei exhibit highly linear age-dependent mHtt DNA repeat expansion (i.e. repeat migration), with modal-CAG increasing at +8.8 repeats/month (R^2^=0.98). This linear rate is reduced to 2.3 and 0.3 repeats/month in Q140 with Msh3 heterozygous and homozygous alleles, respectively. Our study defines somatic *Htt* CAG-repeat thresholds below which there are no detectable mHtt nuclear or neuropil aggregates. Mild transcriptionopathy can still occur in Q140 mice with stabilized *Htt* 140-CAG repeats, but the majority of transcriptomic changes are due to somatic repeat expansion. Our analysis reveals 479 genes with expression levels highly correlated with modal-CAG length in MSNs. Thus, our study mechanistically connects HD GWAS genes to selective neuronal vulnerability in HD, in which *Msh3* and *Pms1* set the linear rate of neuronal mHtt CAG-repeat migration to drive repeat-length dependent pathogenesis; and provides a preclinical platform for targeting these genes for HD suppression across brain regions.

**One Sentence Summary:** Msh3 and Pms1 are genetic drivers of sequential striatal and cortical pathogenesis in Q140 mice by mediating selective CAG-repeat migration in HD vulnerable neurons.

## INTRODUCTION

Huntington’s disease (HD) is a dominantly inherited neurodegenerative disorder with mostly adult onset, relentless progression, and currently no effective disease-modifying therapies^1^. HD is caused by CAG-repeat expansion, encoding an elongated polyglutamine (polyQ) repeat, in exon 1 of *Huntingtin* (*HTT*)^2^. HD patients have CAG alleles ranging from 36 to over 120, and the length of the repeat is inversely correlated with the age at motor disease diagnosis^1^. HD neuropathology is characterized by degeneration of striatal medium spiny neurons (MSNs) and to a lesser extent that of cortical pyramidal neurons (CPNs), accompanied by astrogliosis^3^. A pathognomonic hallmark of HD is the accumulation of mutant HTT (mHTT) protein aggregates in the nuclei and cytoplasm of affected neurons^4^. The exact roles of such aggregates in HD remains unclear^5^.

Genetic mouse models of HD, including an allelic series of murine Huntingtin (*Htt*) knockin (KI) mice^6,7^ with increasing CAG-repeats or human *mHTT* BAC transgenic mice with long uninterrupted CAG-repeats^8^, have shown a triad of striatum-selective molecular and neuropathological phenotypes^1^, including somatic mutant CAG-repeat expansion, transcriptomic dysregulation (termed transcriptionopathy), and mHTT protein nuclear inclusions (NIs) and neuropil aggregates (NAs)^6,8–10^. HD mouse models do not show the overt neuronal loss found in HD^11^, and hence are considered models of early neuronal dysfunction prior to neuronal loss in HD^7^. Currently, it is unclear how *mHTT* CAG-repeat expansion leads to the selective neuronal pathogenesis in either striatal or cortical neurons in HD.

Genome Wide Association Studies (GWAS) have identified 10 genomic loci that significantly modify the timing of HD onset^12,13^ or progression^14^. Strikingly, HD GWAS loci harbor 6 DNA repair genes, including 4 mismatch repair (MMR) pathway genes (*MSH3, MLH1, PMS1* and *PMS2*) and two other genes (*FAN1* and *LIG1*) that may have functional roles in MMR^15,16^. The MMR genes uncovered by the HD GWAS encode components of four evolutionarily conserved MMR protein complexes: MutSβ (MSH2-MSH3), MutLα (MLH1 and PMS2), MutLβ (MLH1 and PMS1) and MutLγ (MLH1-MLH3)^15,17–19^. The MutSβ complex recognizes 2-10 base pair DNA insertion-deletion loops (IDL) as well as repeat-containing extrusions (Fig. 1A), a function that is distinct from MutSα (MSH2-MSH6) that recognizes single mismatch as well as 1-2bp IDLs^20^. Upon recognition of DNA mismatch, the MutSβ complex recruits one of the three MutL complexes to form quaternary complexes^21^. The endonuclease activity of MutLα resects the DNA to allow for repairing mismatched DNA. MutLγ also has endonuclease activity and a known function in meiosis; however, the function of MutLβ, which has no known endonuclease activity, remains unclear^17^. To date, biochemical and genetic analyses have shown that multiple HD GWAS/DNA repair proteins (MSH3, MSH2, MLH1, MLH3 and FAN1) play critical roles in repairing CAG extrusions in vitro^22^ and mutants of their murine homolog genes modifying somatic CAG repeat instability in brain tissues of HD mice^23^. Thus, a major current mechanistic model suggests that GWAS/DNA repair genes accelerate or slow *mHTT* somatic repeat expansion to influence disease onset or progression^24^. However, such a model leaves many unanswered questions. What are the exact mechanistic roles of distinct MMR complex genes (e.g., *PMS2* vs *PMS1*) in HD pathogenesis? Can targeting HD GWAS/MMR genes confer therapeutic benefit beyond consequences for CAG repeat expansion? And if so, what are the magnitudes of disease modifying effects, and can it affect diseases in brain regions beyond the striatum? What are the kinetics of somatic CAG expansion, and how the expanded CAG repeats are translated into neuronal toxicities? And finally, what are the long-term benefits and/or potential harm of targeting key GWAS/MMR genes in the context of HD?

**Figure 1.**
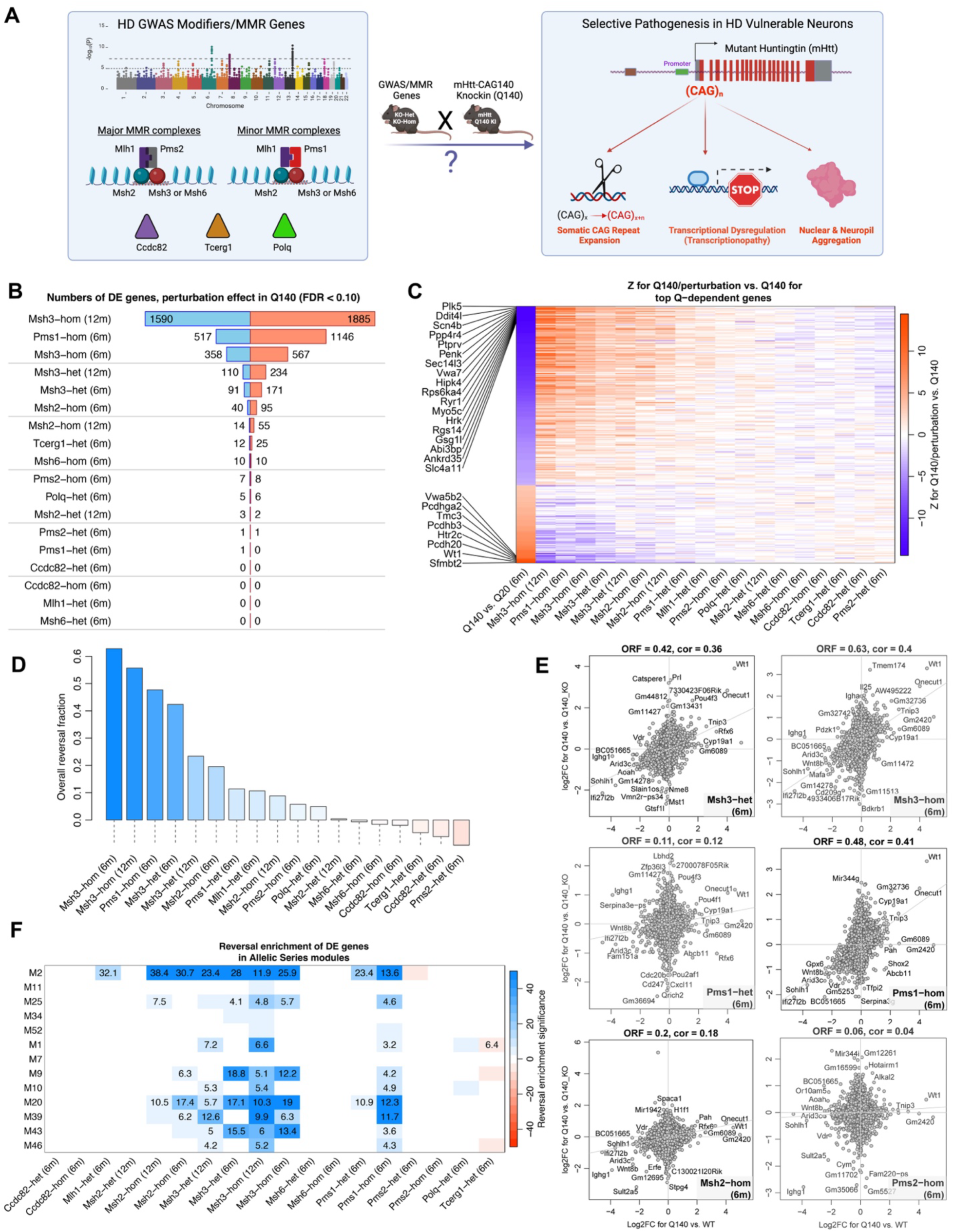
Striatal RNA-seq data from 6- and 12-month-old Q140/GWAS KO mice. (A) Schematic overview of our study. (B) Numbers of genes significantly DE in Q140/KO genotypes vs. Q140 at FDR<0.1. Blue and red bars represent down- and upregulated genes, respectively. (C) For genes DE in our common analysis of Q140 vs. WT at 6m with fold change|>1.5 and meta-average-based FDR<0.05, the heatmap shows meta-average Z statistics for Q140 vs. WT at 6 months and Z statistics for Q140/KO vs. Q140 for the indicated perturbations in the rest of the columns. Each row corresponds to a gene. (D) Barplot of overall reversal fractions (Methods) ordered in decreasing order. (E) Scatterplots of log_2_FC for Q140 vs. Q140/KO (y-axis) against reference Q140 vs. WT (x-axis). Each title indicates the regression coefficient obtained from a linear model regressing y on x. Axis lines and the regression line are shown in grey. (F) Reversal enrichment of DE genes in 13 CAG length-dependent Allelic Series modules (see Methods). Heatmap color indicates “signed” hypergeometric enrichment p-value of genes down- or upregulated in Q140/KO vs. Q140 in each of the modules. Specifically, red (negative) and blue (positive) colors indicate exacerbation and reversal, respectively. Numbers indicate enrichment ratio (observed counts divided by expected counts of overlap genes) for cells with p<10^−4^.

To begin addressing such questions in an in vivo HD model system, we evaluated knockout (KO) alleles for 9 HD GWAS/MMR genes for their modification of a triad of pathological phenotypes that are shared among HD mouse models with long, uninterrupted CAG repeats in mHtt^8^: mHtt CAG-repeat instability, genome-wide transcriptomic dysregulation, and mHTT nuclear aggregation. We also evaluated key disease-suppression mutants for their mechanistic roles in driving neuronal-selective mHtt CAG-repeat expansion and sequential striatal and cortical pathogenesis in a murine Htt knocking mouse model of HD (Q140) with human mHTT Exon1 carrying about 140 CAG repeats replacing murine Htt exon 1^10^.

## RESULTS

### GWAS gene KO mutants reverse striatal transcriptional dysregulation in Q140 KI mice

We tested the hypothesis that the HD GWAS genes may drive transcriptomic dysregulation in Q140 mHtt KI mice^6^. We selected targeted knockout (KO) mutant mice for HD GWAS gene (*Msh3, Mlh1, Pms1, Pms2, Ccdc82, Tcerg1*), and other canonical MMR genes (*Msh2. Msh6*) and a gene (*Polq*) with borderline GWA signal and implicated in MMR^25^. Collectively, we refer to this group of genes as HD GWAS/MMR genes. Seven KO lines are well-characterized null alleles from public repositories, and one allele (Ccdc82 KO) is a CRISPR/Cas9 KO strain generated for this study (see Methods). We crossed these KO mouse lines with Q140 KI/+ mice to accumulate 8 per genotype (sex balanced, Fig. 1A) for all possible genotypes, including, when possible, both homozygous (hom) and heterozygous (het) KO alleles. For *Msh2*, a Cre/LoxP conditional allele was used to avoid early cancer^26^, and MSN-selective Rgs9-Cre^27^ was used to selectively delete *Msh2* in MSNs (referred to as Msh2-hom or Msh2-het). We aged these mice to 6m and dissected striatal tissues for bulk RNA-seq^6,28^. KO lines for *Msh3* and *Msh2*, with previously established effects on CAG instability^23,29^, were also aged to 12m for RNA-seq studies. Besides the striatal tissues, motor-sensory cortex tissues were also dissected from each mouse for RNA-seq. In total we obtained 831 striatal and cortical bulk RNA-seq samples for this study (Fig. S1, Table S1; see Methods).

In the striatum, Q140 vs WT consistently show >5000 differentially expressed genes (DEGs) at 6m in all our cohorts (Table S1). The effects of each GWAS/MMR KO on Q140 transcriptome were assessed by several measures, including the number of DEGs (Fig. 1B, 1C), a transcriptome-wide relationship of log2 fold change (called overall reversal fraction, Fig. 1D, 1E; Methods), and correlation of transcriptome-wide DEG Z statistics (reversal Z-statistics correlation; Fig. S2). Both analyses use Q140 vs WT on the x-axis and Q140 vs Q140/KO perturbation in the y-axis; a positive correlation represents a reversal of Q140 transcriptionopathy by the KO allele while a negative correlation means exacerbation. Together, these analyses reveal that Msh3-hom and Pms1-hom show robust reversal of Q140 striatal transcriptomes at 6m of age with 925 and 1663 significant DEGs (FDR<0.1) between Q140 vs Q140/KO, respectively (Fig. 1B-1E; Fig. S2, S3). Two other KO alleles (Msh3-het and Msh2-hom) show moderate reversal with 262 and 135 DEGs, respectively. Importantly, the transcriptomic rescuing effects of Msh3 in Q140 are not only sustained but also appear to be enhanced at 12m of age with 3475 DEGs for Msh3-hom and 344 DEGs for Msh3-het (Fig. 1B-1E; Fig. S2, S3). Interestingly, the KO alleles alone vs WT for majority of GWAS/MMR genes show minimal transcriptomic effects (<25 DEGs), except for Pms1-hom with 1431 DEGs (Fig. S1A). However, the transcriptomic effects of Pms1-hom in WT or Q140 background are not concordant (Fig. S1B). Our analysis also reveals that our two parameters for ranking transcriptomic reversal are highly correlated with each other (Fig. S3B). Ranking the KO alleles based on their overall rescue fractions, four top perturbations (rescue fractions >0.4) include Msh3-hom at 6m and 12m, Pms1-hom (6m), and Msh3-het (6m); and two moderate rescue (rescue fractions around 0.2) include Msh3-het (12m) and Msh2-hom (6m). Lastly, since Mlh1-hom mice show premature cancer by 5m of age, we analyzed its transcriptomic effects in Q140 at 4m, as the latter already show 804 DEGs (Fig. S4). We found Mlh1-hom shows an overall moderate rescue effect with 11 DEGs and reversal fraction of 0.37 (Fig. S4). Importantly, KO alleles for other HD GWAS/MMR genes (Msh6, Pms2, Tcerg1, Ccdc82, and Polq), elicit 37 or fewer DEGs in Q140 mice, suggesting minimal transcriptomic modifier effects (Fig. 1B-1E; Fig. S2, S3).

We next addressed whether the GWAS gene KOs have similar transcriptomic modifier effects in the motor/sensory cortices of Q140 mice. First, we found the transcriptomic effects of Q140 vs WT at 6m are weak and somewhat variable as in each cohort we found between 4 and 81 DEGs (Fig. S5A, Table S2). Second, in Q140 background, the KO mice in the cortex only yielded 0-25 DEGs for the most KO mice, and only Pms1-hom induced 142 DEGs (Fig. S5B). Moreover, neither the heatmap of DEGs (Fig. S5C) nor the reversal Z-statistics correlation (Fig. S5D) show significant transcriptome-wide changes in the motor-sensory cortical gene expression for the KO alleles in Q140 mice.

Together, our transcriptomic modifier screen of 9 HD GWAS/MMR genes shows a subset of them can robustly and sustainably reverse the striatum-selective transcriptionopathy in Q140 KI mice. Our genetic analyses uncover the essential roles for MutSβ and MutLβ specific genes (i.e., *Msh3* and *Pms1*), but not those specific to MutSα and MutLα (i.e., *Msh6* and *Pms2*), in this disease process.

### Single-nuclei transcriptomics reveals MSN-selective effects of mHTT and correction by *Msh3* mutants

To examine striatal cell-type-specific transcriptional defects and their modification by *Msh3* gene-dosage reduction, we obtained 160,512 high-quality single-nuclei transcriptomes from all 6 genotypes in the Q140/Msh3 KO cross at 12m of age (n=4 animals per genotype, sex-balanced; Fig. S6A). Our clustering pipeline (see Methods) identified 12 cell clusters with strong enrichment of known striatal neuronal and non-neuronal cell type markers (Fig. 2A-2B; Fig. S6; Table S3)^30,31^. The largest two neuronal clusters are annotated as D1-MSN (C1, 55,886 nuclei) and D2-MSN (C2, 44,282 nuclei) (Fig. S6), the two major MSN types that are the most vulnerable in HD (*2*, *4*). Other major cell clusters (Fig. 2A, 2B) include oligodendrocyte (C3), astrocyte (C4), “eccentric” D1-MSN/Tpbg subtype (C5)^30,31^, microglia (C6), oligodendrocyte precursor (OPC, C7), interneurons of various types (C8, C10, C11) as well as mural and endothelial cells (C9) and ependymal cells (C12). In our dataset, the two MSN types constitute 92% of the total striatal neurons and 62% of the total striatal cells, a finding consistent with previous studies^31^ (Fig. S6). Furthermore, the distribution of sequenced nuclei across all six genotypes is roughly even overall (15-19% per genotype) and is in most cellular clusters (Fig. S6). However, we found an overrepresentation of nuclei from the Q140 mice in C3_Oligo and C6_microglia clusters (Fig. S6), which is reminiscent of HD pathology^32^ but requires further validation.

**Figure 2.**
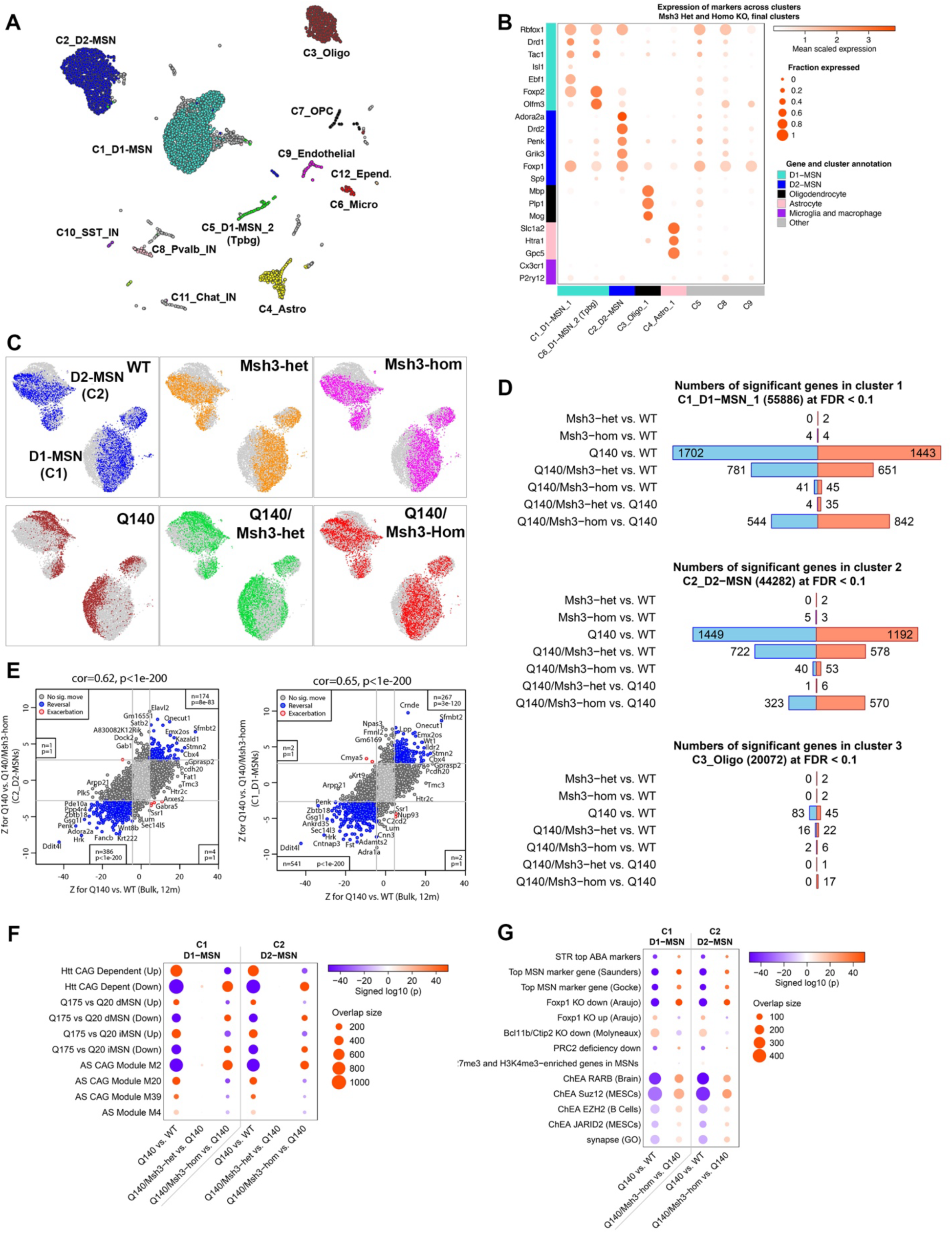
Single nucleus RNA-seq of 12m striata from a Q140/Msh3 KO cross. (A) UMAP representation of snRNA-seq data from WT, Msh3-het, Msh3-hom, Q140, Q140/Msh3-het, Q140/Msh3-hom mouse striatum. Clusters are numbered and labeled by the cell type whose markers show strongest (lowest p-value) enrichment in cluster markers. (B) Expression of selected marker genes (rows) across large clusters. Color intensity indicates mean normalized expression while circle size represents proportion of cells with non-zero expression. (C) UMAP region with clusters C1_D1-MSN and C2_D2-MSN with cells from each of the 6 genotypes in clusters C1_D1-MSN and C2_D2-MSN shown in a distinct color. The color density reflects relative density of cells from the genotype compared to all other cells. (D) Numbers of genes DE at FDR<0.1 in pseudobulk analysis of clusters C1_D1-MSN_1, C2_D2-MSN_ and C3_Oligo. Blue and red bars indicate down- and up-regulated genes, respectively. (E) Reversal scatterplots for Msh3-hom perturbation in clusters C1_D1-MSN_1 and C2_D2-MSN. Each shows Z for Q140 vs. Q140/Msh3-hom (y axis) against Z for Q140 vs. WT in a common analysis of 12-month bulk RNA-seq data from this study. Grey lines indicate approximate locations of FDR=0.1 thresholds. Blue and red color indicate reversed and exacerbated genes at FDR<0.1, respectively. Insets give numbers of genes in each of the corner regions (i.e., genes that pass FDR<0.1 in both tests) and the corresponding hypergeometric p-value. Overall transcriptome-wide correlation (referred to as reversal correlation) and the corresponding nominal p-value are indicated in the title of each plot. (F) Enrichment of DE genes in selected mHtt dependent sets. (G) Enrichment of DE genes in selected cell type marker and TF target sets.

The Uniform Manifold Approximation and Projection (UMAP) embedding reveals a robust segregation of WT cells from Q140 cells in D1-MSN and D2-MSN clusters but not the other cell clusters (Fig. 2C, Fig. S7A). Interestingly, while the cells from WT and Q140 genotypes are largely segregated within the D1- and D2-MSN clusters, Q140/Msh3-het cells are shifted towards WT territory while Q140/Msh3-hom are largely located in WT territory in both MSN clusters (Fig. 3C). To quantitatively evaluate the distribution of cells based on genotypes in each cell cluster, we calculated for each cell the proportion of WT cells among its 30 nearest neighbors (denoted pWT-NN; see Methods). UMAP visualization of pWT-NN shows again a clear separation of areas with high vs low values in C1 and C2 but not in C3, C4 or other clusters (Fig. S7B; data not shown). Statistical testing using pWT-NN aggregated by animal in each genotype found that D1- and D2-MSN clusters (but not oligodendrocyte or astrocyte clusters) show significant differences in pWT-NN across genotypes (Fig. S7C). In both MSN clusters, the WT, Msh3-het and -hom MSNs have >90% WT neighbors, while Q140 MSNs have only about 10% WT neighbors. In Q140/Msh3-het and Q140/Msh3-hom, there is a gene-dosage dependent, significant increase in pWT-NN with the latter approaching about 60%. Thus, *Msh3*-deficiency in Q140 appears to substantially move the dysregulated transcriptome in D1- and D2-MSNs towards those of WT controls.

**Figure 3.**
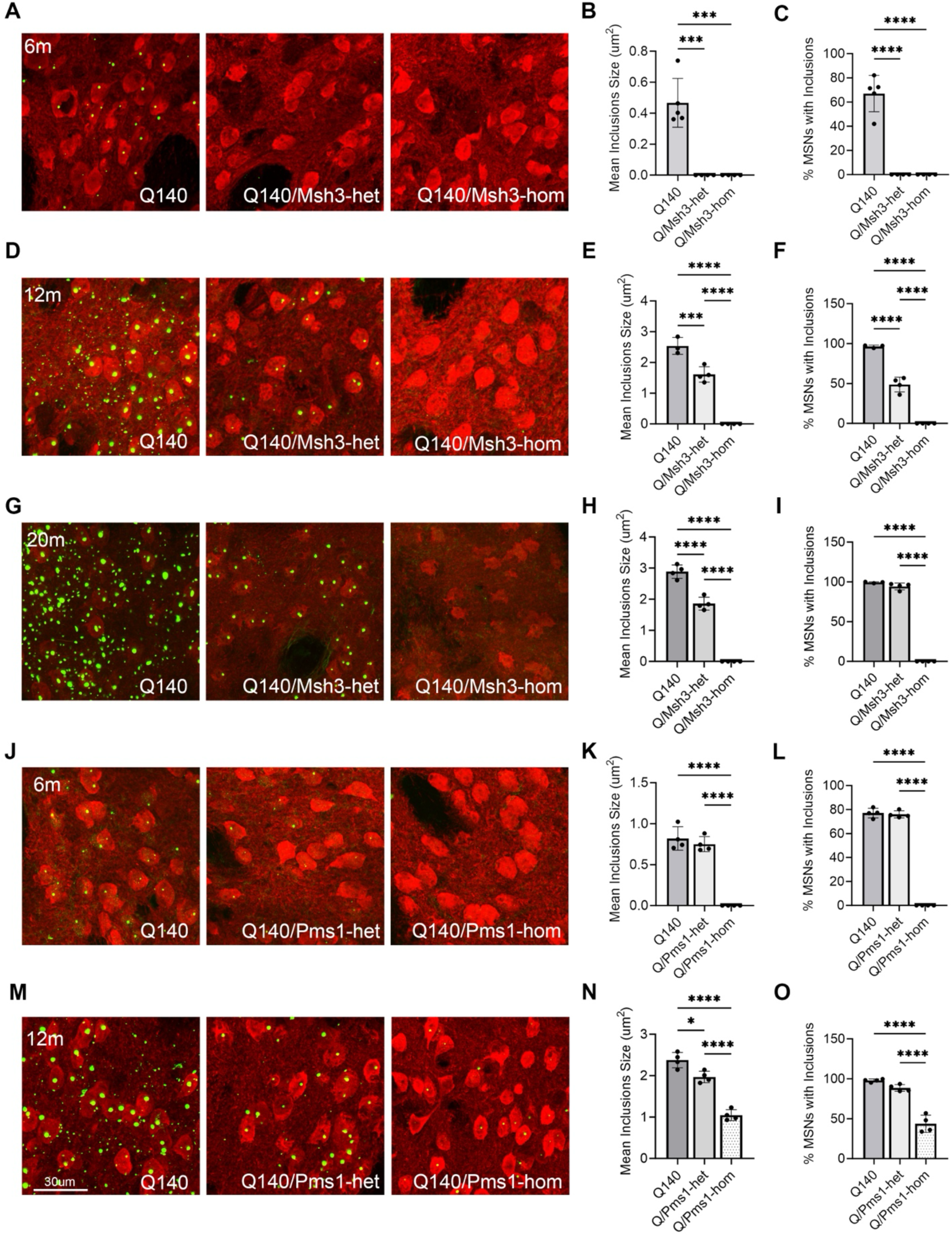
Perturbations rescue mHTT nuclear aggregation in Q140 striata. (A) Representative images of PHP1-stained mHtt aggregates in striata of 6m Msh3/Q140 cohort. Coronal striatal sections (30µm thickness) were double stained with anti-Darpp32 (red) and anti-mHtt (PHP1, green) antibodies. (B) The average size of nuclear inclusion (NI) in striata of 6m Msh3/Q140 cohort. Data are shown as mean ± SEM, n = 3-6 for each age groups. Data analysis by one-way ANOVA with Tukey’s multiple comparison tests: * p<0.05, ** p<0.01, *** p<0.001, **** p<0.0001. (C) The percentage of medium spiny neurons (MSNs) with NIs in striata of 6m Msh3/Q140 cohort. Data presentation is the same to (B). (D) Representative images of PHP1-stained mHtt aggregates in striata of 12m Msh3/Q140 cohort. (E) The average size of NIs in striata of 12m Msh3/Q140 cohort. (F) The percentage of MSNs with NIs in striata of 12m Msh3/Q140 cohort. (G) Representative images of PHP1-stained mHtt aggregates in striata of 20m Msh3/Q140 cohort. (H) The average size of NIs in striata of 20m Msh3/Q140 cohort. (I) The percentage of MSNs with NIs in striata of 20m Msh3/Q140 cohort. (J) Representative images of PHP1-stained mHtt aggregates in striata of 6m Pms1/Q140 cohort. (K) The average size of NIs in striata of 6m Pms1/Q140 cohort. (L) The percentage of MSNs with NIs in striata of 6m Pms1/Q140 cohort. (M) Representative images of PHP1-stained mHtt aggregates in striata of 12m Pms1/Q140 cohort. (M) The average size of NIs in striata of 12m Pms1/Q140 cohort. (O) The percentage of MSNs with NIs in striata of 12m Pms1/Q140 cohort.

We next carried out pseudobulk (single-cell data aggregated by animal) DEG analyses among the genotypes within each cell cluster^33,34^ to avoid potential overestimation of DEGs (Fig. 2D; Fig. S8; Table S3; Methods). These conservative but high confidence analyses reveal 3145 and 2641 DEGs respectively (FDR<0.1; Fig. 2D) in D1- and D2-MSN clusters between Q140 and WT, and they are highly correlated with each other across C1- and C2-clusters (Fig. S8). Impressively, Msh3-het mutation reduces the DEGs between Q140/Msh3-het and WT to 1432 (D1-MSNs) and 1300 (D2-MSNs) respectively, and Msh3-hom further normalize the transcriptome to only 86 and 93 DEGs. The latter represents a 97% reduction of the number of DEGs compared to Q140 vs WT in MSN clusters. Among other cell types in Q140 background, the oligodendrocyte cluster shows 127 DEGs while very few DEGs are found in other cell clusters (Fig. 2D, Fig. S8). In the oligodendrocytes, the DEGs between Q140/Msh3-het or -hom and WT also reduced substantially to 38 and 8, respectively (Fig. 2D). The robust transcriptomic reversal by Msh3-hom in Q140 in the two MSN clusters can also be illustrated by strongly positive transcriptome-wide Z statistics correlation (0.65 and 0.62 in D1- and D2-MSN clusters, respectively) for Q140 vs. Q140/Msh3-hom against those for Q140 vs. WT from 12m bulk RNA-seq (Fig. 2E). Enrichment analyses show Q140/Msh3-hom significantly reverse previously established HD transcriptomic signatures^6,35^, including *mHtt* CAG-length dependent genes and gene coexpression networks in the striatum (Fig. 2F), downregulated MSN identity genes, synaptic genes, and genes regulated or bound by transcription factors (*Foxp1*, *Bcl11b*/*Ctip2*, *Rarb*) or chromatin factors related to PRC2 (Fig. 2G).

Importantly, in the WT background, we did not detect strong transcriptomic effects of Msh3-het or Msh3-hom at 12m (<10 DEGs per cluster; Fig. 2D). Moreover, in either WT or Q140 backgrounds, there is a lack of enrichment of gene signatures associated with disease-associated astrocytes or microglial activation (Fig. S8; Table S3), indicating a lack of overt neuropathology or glial activation.

Together, our snRNA-seq analysis reveals that striatal transcriptionopathy is highly selective to D1- and D2-MSN clusters in 12m Q140 mice, and *Msh3*-deficiency exerts gene-dosage-dependent prevention of such transcriptional dysregulation without eliciting neuronal toxicity or neuroinflammatory gene signatures.

### MMR genes are genetic drivers of striatal MSN-selective mHtt aggregation in Q140 mice

We next examined whether genetic manipulation of HD GWAS DNA repair genes could affect mHTT aggregation, a pathognomonic hallmark of HD^4,8,10,36^. We employed mHTT antibodies PHP1^37^ and EM48^36^, which are commonly used to detect nuclear inclusions (NIs) and neuropil aggregates (NAs) in HD mouse and patient brains. The PHP1^+^ mHtt NIs are absent in 4m Q140 mice and never present in WT mice at any age (data not shown). In 6m Q140, about 60% of DARPP32^+^ MSNs show NIs with average size of 0.4 μm^2^ (Fig. 3A-3C). At 12m and 20m of ages, the striatal aggregates in Q140 show marked progression with NIs occupying nearly 100% of MSNs and their sizes increasing to >2.5 μm^2^ (Fig. 3A-3G). Moreover, the NAs emerge in 12m Q140 striatum and become more numerous at 20m of age (Fig. 3D-3G, Fig. S9), but are almost absent in Q140 at 6m or younger ages (Fig. 3A; data not shown). In Q140/Msh3-het mice, we found no mHtt aggregation at 6m of age; at 12m, mHtt NIs are present only in about 50% MSNs and significantly reduced in sizes to about 1.5um^2^; while at 20m, Q140/Msh3-het still show predominantly NIs in about 100% of MSNs (Fig. 3D-3I). Remarkably, in Q140/Msh3-hom mice, we did not detect any mHtt aggregates (neither NIs nor NAs) in the striatum at 6m, 12m, or 20m of age (Fig. 3A-3I; Fig. S9). We independently verified the 12m Q140/Msh3-hom results using the EM48 antibody at 6 and 12 m (Fig. S10). Next, we confirmed that the reduction or lack of mHtt aggregates in Q140/Msh3-hom or het was not due to any changes in the expression of full-length mutant or wildtype Htt protein levels in their striatal tissues (Fig. S11).

The role of *Pms1* in mHtt aggregation was previously unknown. We examined 6m Q140/Pms1-het mice and found that the frequency and sizes of mHtt aggregation in MSNs are comparable to 6m Q140 (Fig. 3J-3L). Remarkably, in the 6m Q140/Pms1-hom mice, we found a complete absence of mHtt aggregates in the MSNs, revealing *Pms1* as a necessary genetic driver of mHtt aggregation in the striatum. The 12m Q140/Pms1-hom striata show only NIs with significant numbers and sizes, which is similar to those found in 6m Q140 (Fig. 3M-3O). Interestingly, 12m Q140/Pms1-het show a modest but significant reduction in NI size but not frequency compared to Q140 (Fig. 3M-3O). Additionally, we found significant mHtt aggregate reduction in Q140/Msh2-hom (but not Msh2-het) at 6m and 12m (Fig. S12), and for 5m Q140/Mlh1-hom mice (the oldest mice without premature cancer; data not shown). However, we did not detect any changes in mHtt aggregation for the other HD GWAS/MMR genes tested in this study (data not shown). Thus, our study identifies *Msh3* and *Pms1* as genetic drivers of striatal MSN-selective mHtt aggregation and reveals that Msh3 deficiency can sustainably prevent mHtt aggregation in the striatum even in advanced ages.

### Late-onset and progressive cortical pathology in Q140 mice dependent on Msh3 gene-dosage

The cortex is the second most affected brain region in HD^38^. The cortical pathology onset happens later than that of the caudate/putamen and is associated with cognitive impairment and psychiatric symptoms^1^. Although MMR genes such as *Msh3* have been implicated in striatal pathogenesis^29,39^, their role in cortical disease remains unknown. Homozygous Q140 mice are known to have cortical mHtt aggregates^10^, but their distribution in heterozygous Q140 KI mice remains unclear. In PHP1 immunostained brains, we found no cortical mHtt NIs or NAs at 6m of age (data not shown). At 12m, NIs emerge in the piriform cortex while there are few aggregates in other cortical regions such as motor-sensory cortices (Fig. 4A-4C). By 20m of age, the aggregate load in piriform cortex increases both in count and size, and smaller aggregates also are present in motor-sensory and other cortical regions (Fig. 4D-4F). Interestingly, deletion of a single copy of Msh3 is sufficient to abolish most cortical mHtt aggregation in both piriform cortex and motor/sensory cortices at 12m, and diminish the aggregate load at 20m of age to only a few aggregates in the piriform cortex (Fig. 4A-4F). In Msh3-hom, mHtt aggregation is not observed in any of the cortical regions at both 12 and 20m, a finding similar to that found in the striatum (Fig. 2A-2I). Thus, late-onset and progressive cortical mHtt aggregation is a feature of Q140 heterozygous KI mice, and such aggregation appears to be more sensitive to Msh3 deficiency than the striatum as it can be nearly fully suppressed by the deletion of a single copy of *Msh3*.

**Figure 4.**
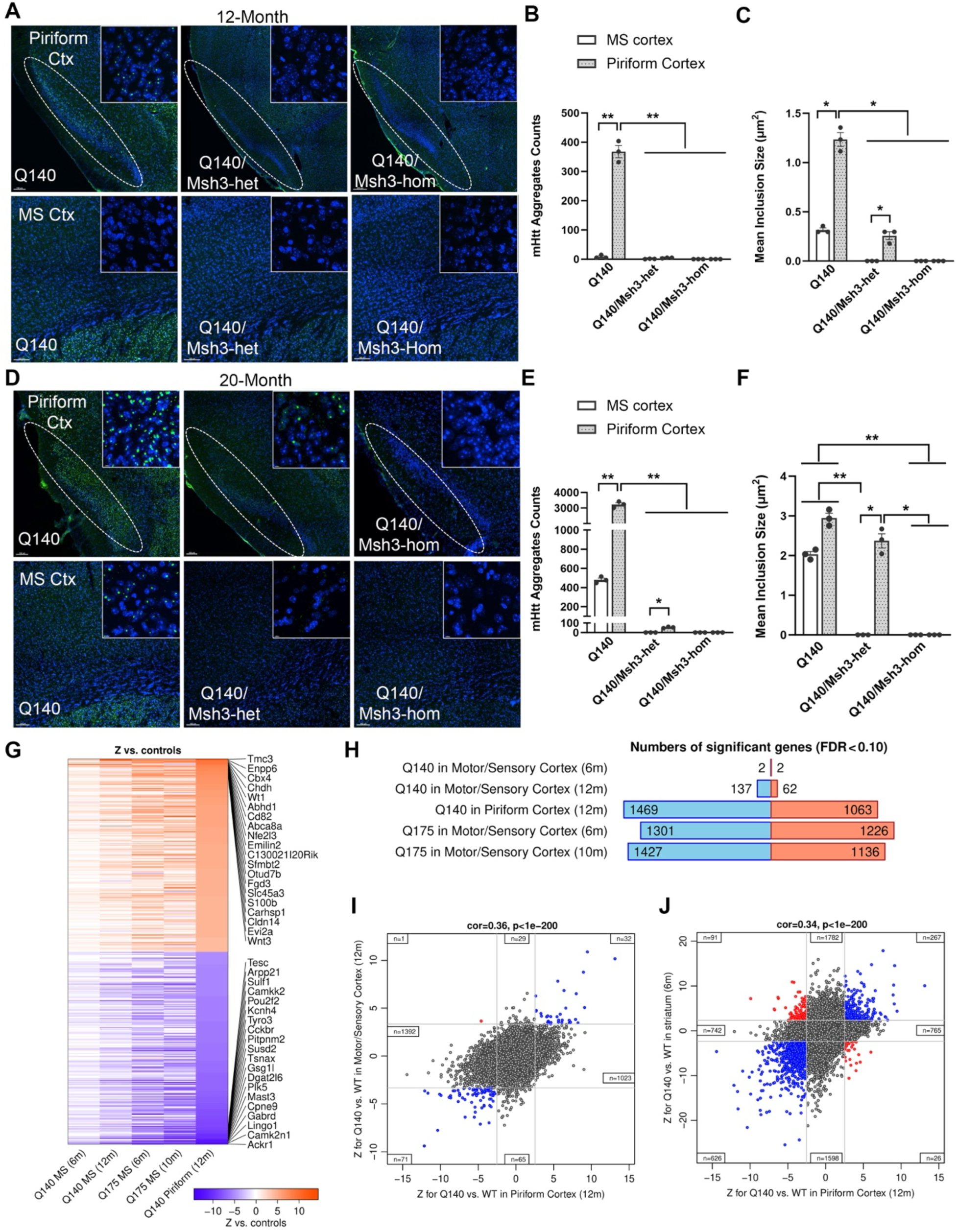
Effect of CAG length expansion and Msh3 KO in mouse piriform cortex. (A) Representative images of PHP1-stained mHtt aggregates in 2 cortical regions – piriform cortex and motor-sensory (MS) cortex of 12m Msh3/Q140 cohort. Coronal striatal sections (30µm thickness) were stained with anti-mHtt (PHP1, green) antibody and DAPI (blue). (B) The total mHtt aggregate counts in 0.08mm^2^ area in the cortices of 12m Msh3/Q140 cohort. Data are shown as mean ± SEM, n = 3-6 for each age groups. Data analysis by one-way ANOVA with Tukey’s multiple comparison tests: * p<0.05, ** p<0.01, *** p<0.001, **** p<0.0001. (C) The average sizes of nuclear inclusion (NI) in the cortices of 12m Msh3/Q140 cohort. (D) Representative images of PHP1-stained mHtt aggregates in 2 cortical regions – piriform cortex and MS cortex of 20m Msh3/Q140 cohort. (E) The total mHtt aggregate counts in 0.08mm^2^ area in the cortices of 20m Msh3/Q140 cohort. (F) The average sizes of NIs in the cortices of 20m Msh3/Q140 cohort. (G) Heatmap representation of Z statistics for Q140 vs. controls for studies indicated in each column labels (MS indicates Motor/Sensory cortex). For Q140 MS (6m), Q140 MS (12m) and Q140 Piriform (12m) the controls are WT mice at the same age, while for Q175 MS at 6 and 10m the controls are Q20 mice at the same age. The heatmap shows top 200 up- and down-regulated genes with largest absolute Z statistics. Each row corresponds to a gene. (H) Numbers of genes DE at FDR<0.1 in the 5 cortex studies included in (G). Blue and red bars indicate down- and up-regulated genes, respectively. (I) Scatterplots of Z statistics for Q140 vs. WT in Motor/Sensory cortex on the y-axis vs. the same DE test in Piriform cortex on the x-axis. Grey lines indicate approximate locations of FDR=0.1 thresholds; insets indicate the numbers of genes in each of the 8 border regions of the plots. Blue and red points represent genes significant at FDR<0.1 in both comparisons that are DE in concordant (blue) and discordant (red) directions. The transcriptome-wide correlation of the Z statistics and its corresponding nominal p-value are shown in the title of each plot. (J) Scatterplots of Z statistics for Q140 vs. WT in common analysis of 6m striatum samples on the y-axis vs. the same DE test in Piriform cortex on the x-axis.

Previous studies in the striatum of BAC-CAG^8^ or Q140 (Fig. 1-2) revealed that germline uninterrupted CAG repeats in mHtt drive transcriptionopathy in addition to mHtt aggregation. We next asked whether the divergence of aggregate load in piriform cortex vs motor-sensory cortex in 12m Q140 mice could also correspond to differential levels of transcriptionopathy. Indeed, RNA-seq reveals 2532 DEGs (FDR<0.1) in piriform cortex while only 199 DEGs in the motor-sensory cortices of 12m Q140 (Fig. 4G, 4H; Table S4). These two cortical regions do share 103 DEGs and an overall moderate Z statistics association (Fig. 4I), suggesting a similar direction of transcriptomic changes with differences in severity. Both the number of DEGs and heatmap analysis reveal the dysregulated genes in Q140 piriform cortices are close to those found in the motor-sensory cortices of Q175 KI mice, a model with about 185 CAG repeats (Fig. 4G, 4H)^40^. Moreover, Z statistics correlations show moderate positive association (R=0.34) with those found in Q140 striatum with 893 shared DEGs, but the majority of DEGs (3380 in the striatum and 1507 in the piriform cortex) in each tissue are not shared (Fig. 4J). Thus, similar to the striatum, progressive cortical mHtt aggregation also co-occurs with robust transcriptional dysregulation in HD mice, and the aggregate pathology is dependent on the presence of two intact copies of *Msh3*.

### A subset of MMR genes drives CAG-repeat instability in striatal and cortical tissues of HD mice

We next asked whether stabilization of somatic mHtt CAG repeats could underlie the correction of transcriptional and aggregate pathologies in Q140 mice by various HD GWAS/MMR gene mutants. To analyze the distribution of mHtt CAG repeats in a tissue, we used an established method^8,41^ involving amplification of the repeat-containing mHtt-exon1 region by PCR, separation of the amplified PCR products by capillary electrophoresis, and subsequent quantification of peaks on both sides of the inherited modal peak (Instability Index Score or IIS) or just those that are larger than the inherited modal peak (Expansion Index Score or EIS). The analysis reveals an expected high IIS and EIS in the striatum and liver of Q140 compared to the cortex (Fig. S13, Table S5). Moreover, KO for *Msh3*, *Msh2*, and *Pms1* all show gene-dosage-dependent reduction of IIS and EIS in the striatum and liver without significant changes in modal CAG repeats (Fig. 5A-5C; Fig. S13). Similarly, we found 4m Mlh1-hom mutants also stabilize the IIS and EIS compared to Q140 (Fig. S13). Importantly, we do not observe any significant effects of *Pms2*, *Msh6*, *Polq*, *Tcerg1* and *Ccdc82* KO alleles on repeat instability in the striatum or liver (Fig. S13, Table S5). Together, we found a robust negative correlation between the transcriptomic reversal (measured by overall rescue factions) the KO lines and the normalized instability scores (measured as IIS in Q140/KO divided by IIS of Q140 in the same cohort; Fig. 5D), demonstrating a robust association between striatal transcriptomic normalization and stabilization of mHtt CAG repeats.

**Figure 5.**
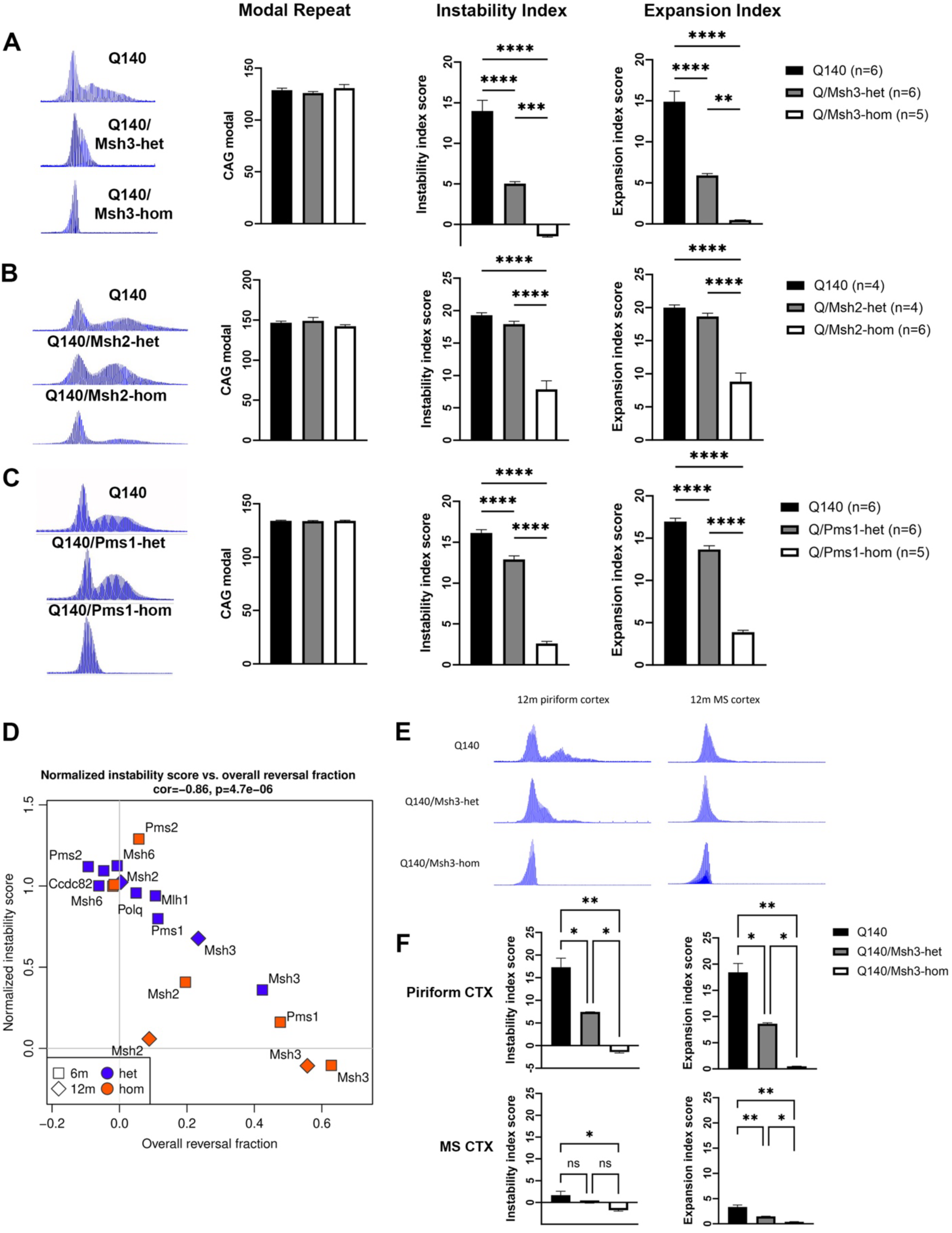
Perturbations rescue somatic CAG instability in the striata and cortices. (A) Representative GeneMapper traces of mHtt CAG repeats and somatic CAG instability indices from striata of 6m Msh3/Q140 cohort. (B) Representative GeneMapper traces of mHtt CAG repeats and somatic CAG instability indices from striata of 6m Msh2/Q140 cohort. (C) Representative GeneMapper traces of mHtt CAG repeats and somatic CAG instability indices from striata of 6m Pms1/Q140 cohort. (D) Scatterplot of normalized instability score (y-axis) vs. overall rescue fraction (x-axis). Ages and zygosities of the perturbations are indicated by point shape and color, respectively. Overall correlation and the corresponding nominal significance (p-value) are indicated in the title of the panel. (E) Representative GeneMapper traces of mHtt CAG repeats in 2 cortical regions – piriform cortex and motor-sensory (MS) cortex of 12m Msh3/Q140 cohort. (F) Somatic CAG instability indices in piriform cortex and MS cortex of 12m Msh3/Q140 cohort.

Since the striatum HD mouse models with long CAG repeats show a triad of mHtt aggregation, transcriptionopathy^6,8^, and CAG repeat expansion (Fig. 1, Fig. 3), we next examined whether the early onset of mHtt aggregation and transcriptionopathy in the piriform cortex compared to motor-sensory cortex could also be associated with enhanced CAG-repeat instability. We isolated piriform and motor-sensory cortical tissues from 12m Q140 and Q140 Msh3 mutants and performed bulk tissue somatic instability analyses. Indeed, we found 12m Q140 mice show much higher CAG repeat instability indices and expansion indices in piriform compared to the motor-sensory cortex. Moreover, in both cortical regions, the IIS and/or EIS are significantly reduced by *Msh3* KO mutants (Fig. 5E, 5F). Together, we conclude the late onset and progressive cortical pathogenesis in Q140 HD mice is also manifested as the pathogenic triad and is driven by Msh3.

### Linear rate of migration of mHtt CAG repeats in striatal MSNs and its dependency on Msh3 gene-dosage

Analyses of CAG-repeat expansion in HD mouse models thus far only show mosaic patterns of repeat expansion at tissue levels, with certain tissues (e.g., striatum and liver) showing higher levels of expansion then others (cortex, cerebellum and tail)^8,41^. However, given such analysis using the whole tissue with mixed cell types, it remains unclear whether the HD vulnerable neurons (e.g., striatal MSNs) still show mosaic patterns of expansion or a higher, more uniform pattern of expansion. Recent studies using postmortem HD brain samples show a subset of neurons, including those striatal and cortical neurons that are vulnerable to degeneration in HD, exhibit a large separation of modal CAG repeat from the inherited mutant alleles^39,42^. However, the human studies have not yet discerned the kinetics of such neuronal-selective CAG-repeat migration and its relationships to GWAS/MMR variants. Here we hypothesize that Q140 mice show MSN-selective mHtt modal CAG repeat migration, and the observed tissue level modal repeat peak at the inherited allele size is due to the non-MSN cell types.

To test this hypothesis, we first used Fluorescent Activated Nuclei Sorting (FANS) with anti-NeuN antibody to purify all the neuronal nuclei from 6m Q140 striata and performed CAG repeat sizing using similar PCR-based method used for tissues (see Methods). Since MSNs constitute about 90% of striatal neurons while about 10% are interneurons, we found that the CAG distributions still have a modal peak at the inherited ∼140 CAG and a second peak representing the somatically expanded CAG alleles (Fig. S14). We next examined striatal mHtt CAG distribution in MSN nuclei purified with FANS using antibody against Bcl11b (or Ctip2), an MSN specific transcription factor^43^. Remarkably, we found the modal CAG repeats (termed *Mod-CAG*) for Q140 MSNs progressively shift right-ward with age at 5 timepoints, from ∼137 repeats at 2m to up to ∼260 repeats at 16m (Figure 6A). Importantly, the distribution of CAG repeats in Q140 MSNs lacks a peak at 140-CAG in brains from 4m and older animals, despite that the PCR-based CAG repeat sizing assay has an inherent bias towards the lower repeat sizes. These results suggest that the CAG repeats in MSNs are expanding at an MSN population scale in a relatively synchronous manner (termed hereafter as *CAG-repeat migration*). Moreover, we used FANS-purified MSNs to show Mod-CAG migration is greatly slowed down in Q140/Msh3-het, further stabilized in Q140/Pms1-hom, and fully stabilized at the inherited allele in Q140/Msh3-hom at 6m, 12m and 20m of age (Fig. 6B-6D).

**Figure 6.**
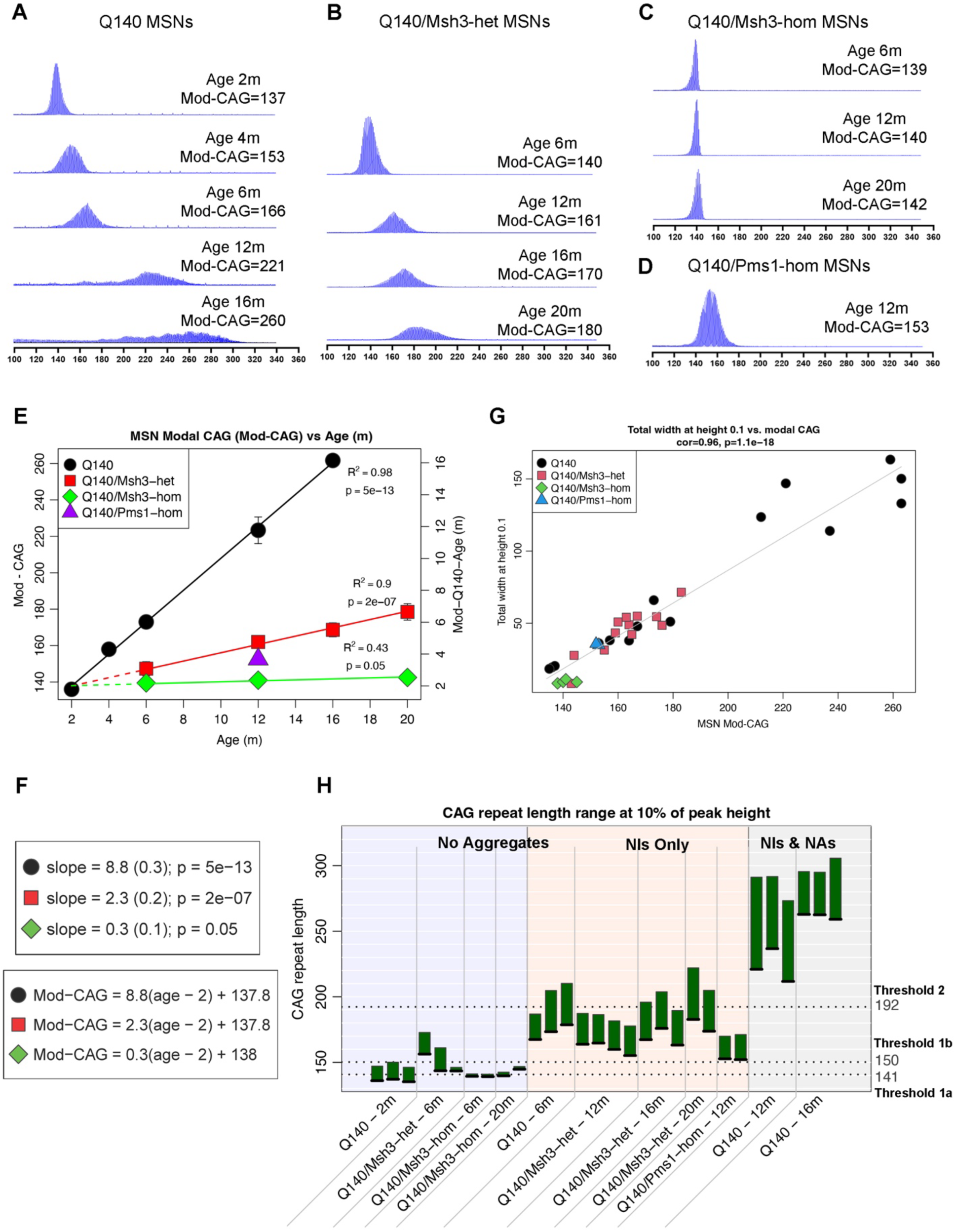
Linear rate of migration of mHtt CAG repeats in striatal MSNs depends on Msh3 and Pms1. (A) Representative GeneMapper traces of mHtt CAG repeats in MSN nuclei purified with FANS using antibody against Bcl11b (or Ctip2) from Q140 striatal tissues at different ages. (B) Representative GeneMapper traces of mHtt CAG repeats in purified MSN nuclei from Q140/Msh3-het striatal tissues at different ages. (C) Representative GeneMapper traces of mHtt CAG repeats in purified MSN nuclei from Q140/Msh3-hom striatal tissues at different ages. (D) Representative GeneMapper traces of mHtt CAG repeats in purified MSN nuclei from 12m Q140/Pms1-hom striatal tissues. (E) Scatterplot of modal CAG lengths (Mod-CAG, y axis) vs. chronological age (x-axis) for Q140 (black circles), Q140/Msh3-het (red squares), Q140/Msh3-hom (green diamonds) and Q140/Pms1-hom (purple triangle). For the first three genotypes, linear model fit lines are shown in the corresponding color. The linear models for Q140/Msh3-het and -hom include the 2m Q140 Mod-CAG lengths; this fact is indicated by extending the corresponding regression lines to 2m using dashed lines. For each of the three linear models, the adjusted R^2^ and the significance of the coefficient (more precisely, the p-value for the null hypothesis that the coefficient is zero) are shown next to the regression lines. Scale on the right side of the plot indicates age (Mod-Q140-Age) predicted using the linear model of age as a function of Mod-CAG in Q140 samples. (F) Linear model coefficients (slopes) and p-values for each of the linear models (regression lines) in panel (E), and formulas relating Mod-CAG to age in each of the three genotypes. The formulas apply only to the age range within each genotype, i.e., between 2 and 16m in Q140 and between 2 and 20 months in Q140/Msh3-het and -hom. (G) Scatterplot of total width at 10% of peak height cs. Mod-CAG. Shape and color of points indicates genotype. (H) Bars indicate the range of CAG repeats between the modal CAG length (black line at the bottom of each bar) to the right CAG peak boundary at 10% of peak height (top of each bar). Each bar corresponds to a single sample (animal). Threshold 1a (lowest dotted line) represents the mean of Mod-CAG lengths in Q140/Msh3-hom at 6 and 20m; threshold 1b represents the mean of the upper CAG lengths in Q140/Msh3-het at 6m and Q140/Msh3-hom at 6 and 20m; and threshold 2 represents the mean of the upper CAG lengths in the “NIs Only” group, i.e., Q140 at 6m, Q140/Msh3-het at 12, 16 and 20m, and Q140/Pms1-hom at 12m.

Given our precise quantitation of MSN Mod-CAG repeats at multiple ages in Q140 and Q140 with MMR mutations, we performed regression analysis to investigate the rates of CAG repeat changes in these various genotypes. We found that a linear model (Fig. 6E) fits exceedingly well for Q140 (adjusted R^2^=0.98) and Q140/Msh3-het (R^2^=0.94), and moderately well for Q140/Msh3-hom (R^2^=0.43). The regression line slope, representing the rate of CAG expansion, is 8.8 repeats/month in Q140 MSNs, which is reduced to 2.3 repeats/month for Q140/Msh3-het, and 0.3 repeats/month for Q140/Msh3-hom (Fig. 6F). Our studies allow for the first time to use simple linear equations (Figure 6E) to accurately predict, based on chronological age, a parameter (i.e. Mod-CAG) associated with MSN-selective CAG expansion in the brain of mammalian HD model. Moreover, for HD mice with genetic (e.g. Msh3 KO) or other perturbations, we can obtain MSN Mod-CAG and use the formula for Q140 MSNs to calculate the equivalent of “Mod-CAG age for Q140” (called Mod-Q140-Age). Thus, a 20m Q140/Msh3-het has a Mod-Q140-Age of about 6m, a 14m deceleration of CAG expansion; and a 20m Q140/Msh3-hom has a Mod-Q140-Age of about 2m, and hence 18m deceleration of CAG expansion (Fig. 6E).

Besides Mod-CAG migration, we noticed that the shape of the MSN CAG repeat distribution in Q140 is also progressively widening with age, suggesting a modest degree of CAG repeat dispersion during mostly synchronous repeat expansion in the MSNs. We confirmed that the MSN Mod-CAG is highly positively correlated with the width of CAG peaks (measured at 0.1 height at the modal repeats) across all four genotypes, i.e., Q140 as well as Q140 with *Msh3* and *Pms1* KO mutants (Figure 6G; Pearson correlation=0.96, p=1.1e-18). The mechanism underlying this pattern of modest CAG repeat dispersion during repeat migration is unclear, but it further demonstrates the quantitative rigor of our MSN-selective CAG repeat measurement and its suitability to study the relationships between CAG-repeat expansion and neuronal pathogenesis in HD mice.

### Mutant Htt aggregation requires CAG-expansion beyond thresholds in striatal MSNs

A striking finding in this study is that Q140 mice fully lacking *Msh3* do not show any mHtt NIs or NAs at 12m and 20m of age, when 100% of MSNs in Q140 with intact *Msh3* show such pathology (Figure 3). This finding is consistent with the hypothesis that aggregation can occur at the inherited mHtt CAG repeat size of 140 and repeat expansion within MSNs is mosaic with a portion of mHtt remaining unexpanded by 12m or 20m of age. Given our finding that mHtt CAG repeats in Q140 MSNs show a relatively synchronous repeat migration with essentially no CAG alleles remains at 140 at ages of 12m and above, the all-or-none mHtt aggregation phenotype in Q140 vs Q140/Msh3-hom alleles can be simply explained by a model in which mHtt aggregation in MSNs, at least those that can be detectable by PHP1 and EM48 antibodies, requires CAG repeat expansion beyond the inherited 140 CAG repeats (Fig. 6H, Threshold 1a). To further refine this idea, we categorized genotypes/ages of the mice in our study into three groups, those without any mHtt NIs or NAs, those with only NIs but not NAs, and those with both NAs and NIs (Fig. 6H). We reasoned that the upper threshold of MSN CAG repeats for transitioning from one category to the next can be estimated using the average of the CAG sizes for all the genotypes within the same category. Since we want to conservatively estimate the threshold below which there is lack of transition (e.g., no aggregates to NI, or NI only to NI plus NA), we used for each group, regardless of genotypes/ages, the upper range of CAG repeat spread at 10% modal heights. With such an analysis, we found that the average repeat length for all genotypes with no aggregates is below 150 CAG (Threshold 1b), and that for those having NIs but lacking any NAs is 192 CAG (Fig. 6H). These estimated thresholds are consistent with other key prior findings in HD mouse models: (i). KI mice with less than 140 inherited CAG alleles (e.g. Q92, Q111) take much longer time to accumulate NIs^44^; (ii). mouse models with stable CAA-CAG repeats encoding 226Q^45^, but not 97Q^46^, show robust NI pathology; and importantly, Q175 mice with inherited repeats (185 CAG) beyond our aggregation threshold of 150 CAG no longer show amelioration of aggregation phenotype with Msh3 KO^47^. Together, our study provides the first robust evidence that somatic huntingtin CAG expansion beyond a threshold is required to elicit mHtt nuclear and neuropil aggregation pathology, a pathognomonic phenotype in HD brains.

### Deconvolution of somatic CAG repeat length and age in eliciting transcriptional dysregulation in HD mice

Transcriptomic analyses of the allelic series KI mice showed that the inherited CAG-length is associated with striatum-selective^6^, and MSN-predominant^35^ transcriptionopathy in HD mice. Here we explore further two questions: (i). Is CAG expansion necessary to induce transcriptional dysregulation in a manner analogous to how it is necessary for mHtt aggregation (e.g. beyond a threshold that is above 140 CAG); and (ii). Is somatic CAG-repeat expansion in MSNs associated with dysregulation of specific genes in a repeat-length dependent but genotype-independent manner? The latter genes could potentially serve as gene expression biomarkers for interventions that can stabilize or contract the somatic HTT CAG repeats in vivo.

To address the first question, we analyzed DEGs between Q140/Msh3-hom mice and WT mice at 6m and 12m as the CAG repeat length of 140 is stable at these ages (Fig. 6C, 6E; Fig. 7A). While Q140 show 4434 and 6116 DEGs (FDR<0.1) compared to WT at 6m and 12m, respectively, Q140/Msh3-hom only shows 103 and 1464 DEGs, respectively, compared to WT. Thus, with stabilized CAG repeats, Q140 mice only showed 1.9% and 19.6% DEGs at 6m and 12m of age, representing a substantial delay in the onset and much lesser severity of this phenotype. However, transcriptome-wide Z statistics correlation for 12m DEGs in the two comparisons still show high positive concordance (R=0.66), suggesting stabilized 140 CAG dysregulate similar genes but to a much lesser degree (Fig. 7B). Thus, we conclude that striatal transcriptional dysregulation does not absolutely require CAG expansion beyond 140 CAGs, but such somatic CAG expansion greatly accelerates and exacerbates the transcriptionopathy.

**Figure 7.**
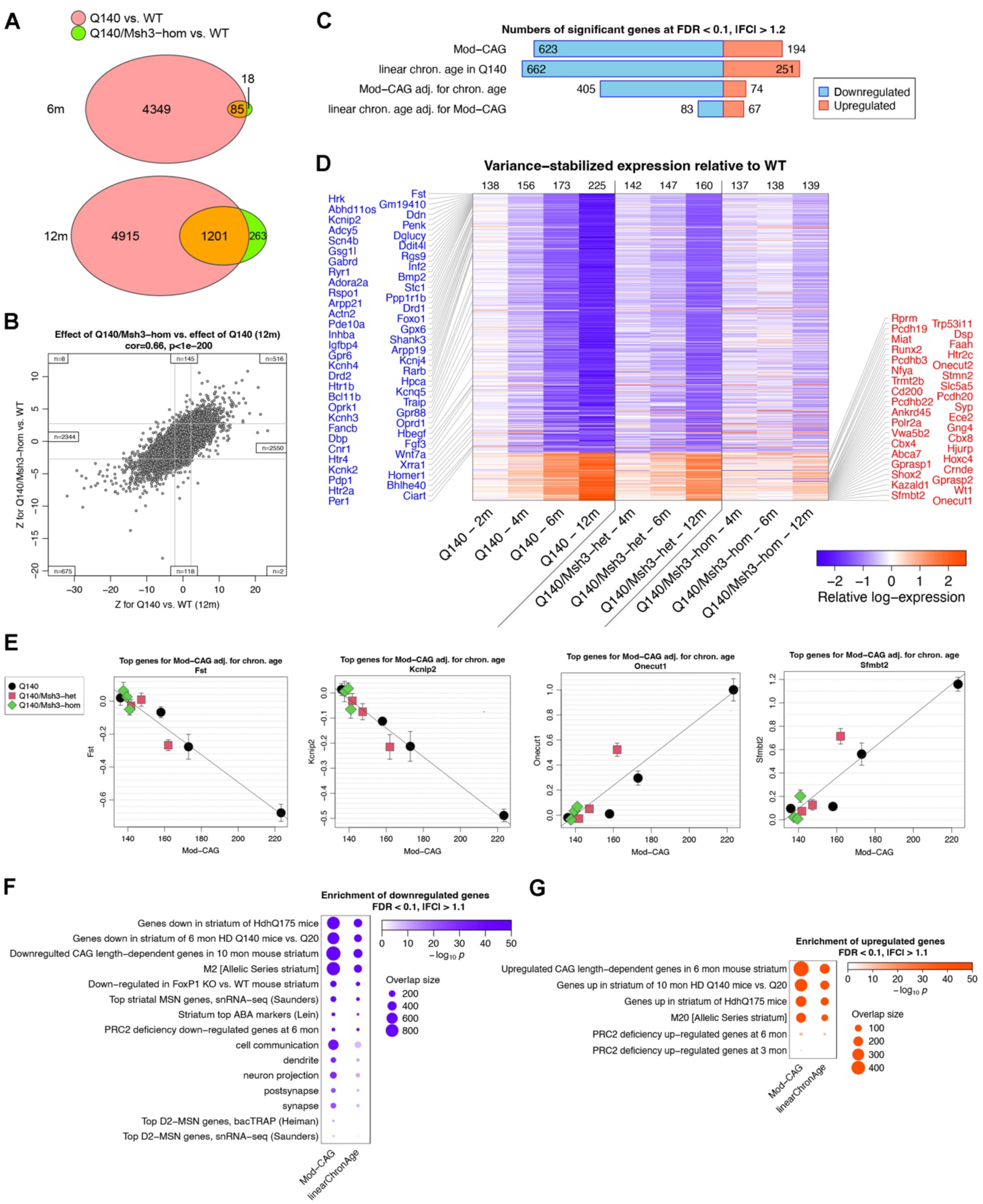
Somatic CAG-repeat-length and age effects HD transcriptional dysregulation. (A) Venn diagrams representing overlaps of genes DE at FDR<0.1 in Q140 vs. WT and Q140/Msh3-hom vs. WT at 6 and 12m. (B) Scatterplot of DE Z statistics for Q140/Msh3-hom vs. WT at 12m (y axis) against meta-average Z for Q140 vs. WT in the common analysis of 12m samples reported in this work. Vertical and horizontal grey lines show approximate location of FDR=0.1 thresholds. Insets report numbers of genes in each of the 8 border regions of the plot. Transcriptome-wide correlation of the Z statistics and the corresponding p-value are shown in the plot title. (C) Bars report numbers of genes associated at FDR<0.1 and |FC|>1.2, where fold change represents changes between 12 and 2m for age tests and between the Q140 Mod-CAG at 12m and at 2m for Mod-CAG tests. The four bars represent tests for linear association of gene expression with Mod-CAG, chronological age in Q140 samples only, Mod-CAG adjusted for chronological age and chronological age adjusted for Mod-CAG. (D) Each column of the heatmap represents the difference of mean expression in the genotype and age indicated below the column and the WT controls at the same age; each row corresponds to a gene. Numbers at the top of each column indicate the predicted modal CAG length for each genotype and age combination. (E) Illustrative scatterplots of genes strongly down- and upregulated with increasing Mod-CAG when adjusted for chronological age. (F) Selected terms illustrating enrichment of genes downregulated with increasing Mod-CAG and linear chronological age at FDR<0.1 and |FC|>1.1. (G) Selected terms illustrating enrichment of genes upregulated with increasing Mod-CAG and linear chronological age at FDR<0.1 and |FC|>1.1.

We next sought to identify genes whose striatum expression levels are highly associated with the Mod-CAG length adjusted for age as well as those that are highly associated with linear age adjusted for Mod-CAG. Our analysis included transcription and either measured or predicted (using our linear models) Mod-CAG data for Q140 (2, 4, 6, 12m) and Q140/Msh3-het and -hom (4, 6, 12m) genotypes. Because each age in our RNA-seq data forms a separate batch, we first performed batch correction designed to equalize the mean values of each gene across WT samples from all ages. Thus, our analysis is assessing the effects of Mod-CAG and linear age on gene expression levels of Q140 or Q140/Msh3 KO mutants relative to their respective WT samples. We identified genes with highly significant (FDR<0.1; FC>1.2) linear association with Mod-CAG (817 genes) or linear age (913 genes; Fig. 7C, Table S6). Importantly, we defined 479 genes significantly associated with Mod-CAG adjusted for linear age and 150 genes significantly associated with linear age adjusted for Mod-CAG (Fig. 7C). At the transcriptome-wide level, the correlation of Z statistics for Mod-CAG adjusted for age with Z for linear age adjusted for Mod-CAG is moderate (R^2^=0.38; Fig. S15), suggesting the effects of Mod-CAG and age differ substantially. Interestingly, genes downregulated with Mod-CAG adjusted for age are far more numerous than upregulated ones (405 vs. 74), while such a bias is not apparent for those associated with linear age (83 and 67). This result suggests somatic MSN Mod-CAG expansion preferentially downregulates gene expression in Q140 striatum.

Genes strongly linearly associated with Mod-CAG adjusted for age are of high interest because they could provide insights into mechanisms underlying somatic expansion-related neuronal toxicities. Heatmap visualization confirms the expression levels of these genes (relative to WT) are highly dependent on Mod-CAG lengths regardless of genotype or age (Fig. 7D). Furthermore, the top Mod-CAG-downregulated (*Fst*, *Kcnip2*) or upregulated (*Onecut1*, *Sfmbt2*) genes show highly linear associations with repeat lengths (Fig. 7E). The genes downregulated with Mod-CAG are enriched in DEGs from HD KI mice (Q140 and Q175), genes in mHtt-downregulated module M2 module from the Allelic Series^6^, markers of striatum MSNs, and genes downregulated in the striatum by FoxP1 KO and PRC2 deficiency (Fig. 7F, 7G). FoxP1 and PRC2 are known to be critical in establishing the MSN identity gene expression, synaptic connectivity, and survival^48,49^. Moreover, the downregulated genes are enriched with GO terms such as synapse/post-synapse, neuronal projection/dendrite, and cell communication. The enrichment terms for genes upregulated with Mod-CAG include genes upregulated HD KI mice, the top mHtt-upregulated gene coexpression module M20^6^, and PRC2-deficiency upregulated genes^49^. Interestingly, the latter group contains multiple neurodevelopmental transcription factors (e.g., *Onecut1*, *Sfmbt2*, *Wt1*, *Cbx4*, *hoxc4*, *Shox2*) that are normally silenced by PRC2-mediated bivalent chromatin and their re-expression can lead to MSN degeneration in PRC2-deficient mice^49^. Additionally, multiple Mod-CAG-downregulated genes are involved in neuronal activity changes in MSNs, e.g., sodium (*Scn4b*) and potassium channels (*Kcnh3*, *Kcnh4*, *Kcnj4*, *Kcnk2*, *Kcnq5*) while others implicate metabolism (*Pdp1*), circadian clock (*Per1*, *Dbp*, *Bhlhe40*), and DNA repair (*Xrra1*, *Rprm*, *Traip*), which all warrant further investigations. Finally, multiple Mod-CAG downregulated genes are known to be involved in hereditary movement disorders, i.e. *HPCA*^50^ and *ANO3*^51^ in familial dystonia, *ADCY5* and *PDE10a* in childhood chorea^52^, which are all aspects of motor deficits in HD. Lastly, *Gsg1l* is a synaptic gene^53^ that is located in an HD GWA significant locus^13^, hence it could be a candidate that connects somatic CAG expansion to synaptic deficits in HD.

Finally, the 479 Mod-CAG associated genes adjusted for linear age provide valuable candidates for gene expression biomarkers for interventions to stabilize or contract CAG expansion in HD MSNs. To this end, there genes on this list that encode proteins with known PET ligands either already tested in HD (*Pde10a*, *Drd1*, *Drd2*)^54^ or yet to be tested (*Htr1b*, *Htr2a*, *Htr4*, *Oprd1*, *Oprk1*). Other genes encode secreted factors (e.g., *Fst*, *Inhba*, *Igfbp4*, *Cx3cl1*, *Fgf3*, *Hbegf*, *Wnt7*) or enzymes (e.g., *Ace*), which could potentially be measured in biofluids such as CSF. Further studies of these genes and their protein products, especially in patients and their biosamples, will be key to advance the identification of molecular or imaging biomarkers for somatic CAG expansion in HD.

### Msh3 mutants reverse neuropathology and locomotor deficits in HD Q140 mice

Synapse loss in the striatal MSNs is a shared feature of HD mouse models and patients^8,55,56^, and our enrichment analysis reveals Q140/Msh3-hom prevents the downregulation of synaptic genes (Fig. 2G). In 12m Q140 striata, we found a significant reduction of striatal postsynaptic marker protein Actn2 in two independent cohorts (Fig. 8A, 8C; Fig. S16). Importantly, both Q140/Msh3-hom and Q140/Msh2-hom, but not their respective Q140/KO-het mice, significantly enhanced Actn2 levels compared to the Q140 controls, and restored them to levels similar to WT (Fig. 8A; Fig. S16).

**Figure 8.**
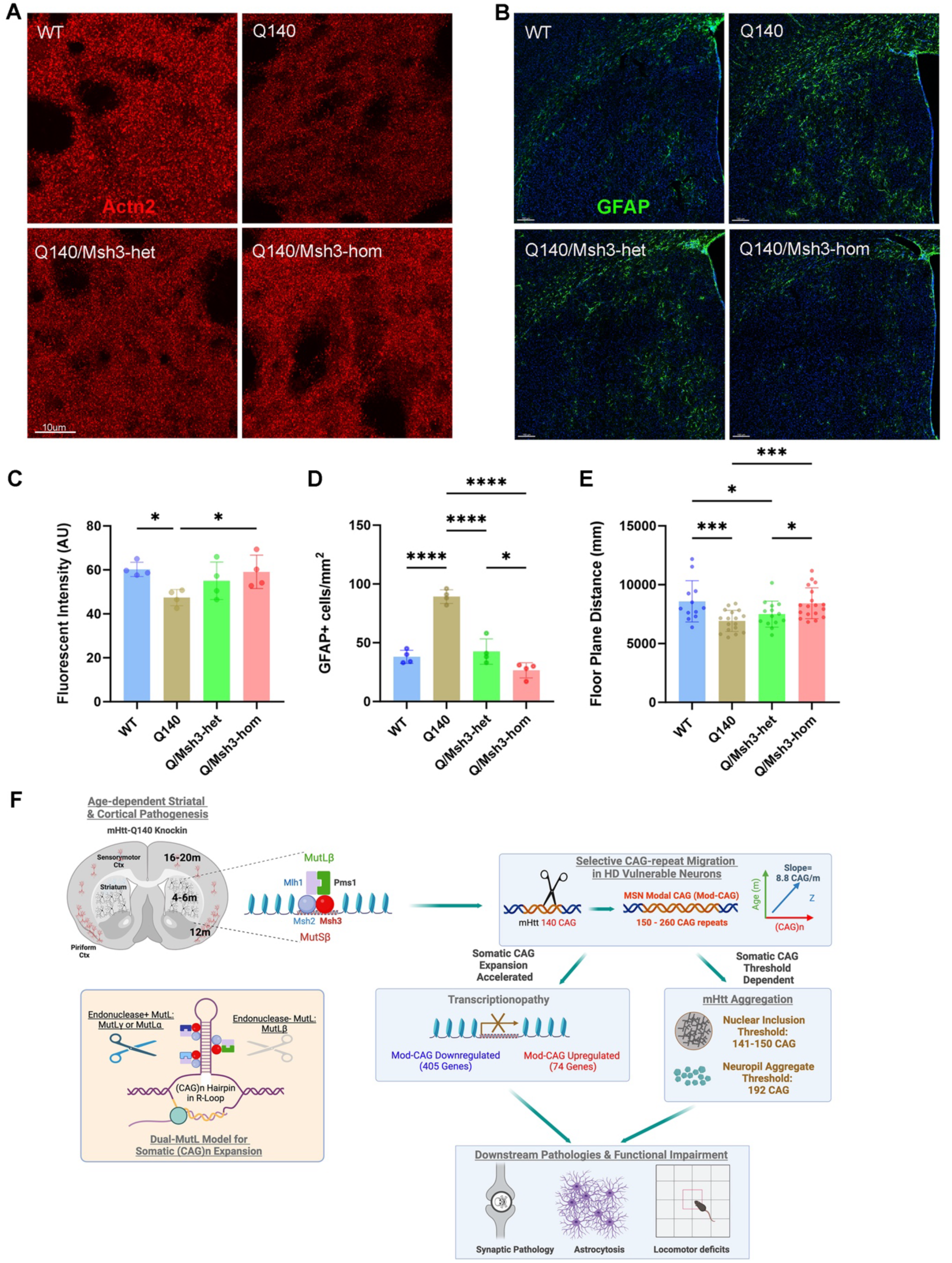
Msh3 mutants reverse neuropathology and locomotor deficits in HD Q140 mice. (A) Representative images of Actn2 (post-synaptic marker) staining in striata of 12m Msh3/Q140 cohorts. Coronal striatal sections (10µm thickness) were stained with anti-Actn2 (red) antibody. (B) Representative images of GFAP (astrocytosis marker) staining in 20m Msh3/Q140 cohorts. Coronal striatal sections (30µm thickness) were stained with anti-GFAP (green) antibody and DAPI (blue). (C) Quantifications of Actn2 intensity in striata of 12m Msh3/Q140 cohorts. Results are shown as mean ± SEM from 4 brains for each group. Data analysis by one-way ANOVA with Tukey’s multiple comparison tests: * p<0.05. (D) Quantifications of GFAP+ cell density in striata of 20m Msh3/Q140 cohorts. Results are shown as mean ± SEM from 4 brains for each group. Data analysis by one-way ANOVA with Tukey’s multiple comparison tests: * p<0.05, **** p<0.0001. (E) Msh3 KO rescued hypo-locomotion behavior in 6m Q140 measured by the Open-field test. Results are shown as mean ± SEM from 12-18 mice for each group. Data analysis by one-way ANOVA with Tukey’s multiple comparison tests: * p<0.05, *** p<0.001. (F) A schematic mechanistic model illustrating that genes encoding the MutSβ-MutLβ complex are genetic drivers of selective neuronal pathogenesis in HD mouse model. These protein complex, working in conjunction with one of the MutSβ--MutL complexes with endonuclease activity (i.e. MutLα or MutLγ), to drive neuronal-selective CAG repeat migration to elicit CAG-length dependent mHtt aggregation, transcriptionpathy and downstream toxicities (e.g. synaptic marker protein loss, astrocytosis, and locomotor deficits.

We next examined whether long-term deficiency of Msh3 in wildtype or Q140 mice at 20m of age focusing on GFAP, a marker of astrocytosis that is upregulated in HD patient caudate/putamen and a key component used in determining HD neuropathological grades^57^. We found that 20m Q140 mice showed a significant upregulation of GFAP compared to WT in the dorsal striatum, mimicking HD neuropathology (Fig. 8B). Impressively, both Q140/Msh3-het and Q140/Msh3-Hom showed significant reduction of GFAP staining in the striatum (Fig. 8B, 8D). The levels of GFAP in Q140/Msh3-hom is not significantly different from those of WT at this age (Fig. 8B, 8D). Thus, long-term Msh3 deficiency not only reduces mHtt aggregation but also shows evidence of reducing astrocytosis in Q140 mice. Additionally, long-term deficiency of Msh3 does not show any evidence of increasing microgliosis compared to WT by Iba1 staining (data not shown), again suggesting Msh3 deficiency does not elicit age-dependent glial cell activation in vivo.

Finally, no prior studies have addressed whether genetic manipulation of an MMR genes such as Msh3 could modify the behavioral deficits in an HD mouse model. To address this important question, we assessed whether *Msh3* deficiency could modify behavioral deficits in Q140. At 6 months, Q140 mice compared to WT controls showed reduced nighttime locomotion in an open field test, which was significantly rescued in Q140/Msh3-hom but not in Q140/Msh3-het mice (Fig. 8E). Together, our study suggests that mice lacking Msh3 show strong evidence of improving neuropathology and behavioral deficits in Q140 mice beyond CAG repeat stabilization and aggregate reduction, and these long-term functional benefits in HD mice have not previously been reported in studies using genetic manipulation of HD GWAS/MMR genes such as *Msh3*.

## DISCUSSION

### Distinct MMR complex genes drive selective neuronal pathogenesis in HD mice

Our study provides strong evidence that mice with KO of four HD GWAS/MMR genes either strongly (*Msh3* and *Pms1*) or moderately (*Msh2* and *Mlh1*) reduce the triad of striatum-selective pathogenesis in HD mice: CAG-repeat instability, mHtt aggregation and transcriptionopathy (Fig. 8F). Our study reveals specific MMR complex genes are crucial to HD pathogenesis in vivo. First, deletion of Msh3 but not Msh6 exerts robust, gene-dosage-dependent amelioration of HD-related phenotypes in Q140, demonstrating MutSβ rather than MutSα as key to disease modification, a finding consistent with the fact that *MSH3* but not *MSH6* is located one of the HD GWAS loci^13^. Our genetic results are in full synergy with biochemical findings that the MutSβ (but not MutSα) specifically recognizes the CAG loop-out substrate to mediate its repair in vitro (Iyer & Pluciennik, 2021). For MutL complexes, current models^16,21^ suggest that the MutLα (MLH1-PMS2) as well as MutLγ (MLH1-MLH3) could be key, as these complexes but not MutLβ (MLH1-PMS1) have endonuclease activities that are postulated to cut the DNA and initiate the repeat expansion process. In this context, our finding that Pms1 but not Pms2 is essential for eliciting the striatal pathogenic triad is a somewhat surprising finding. The yeast Pms1 homolog (Mlh2) is known to be recruited to DNA foci upon induction of DNA damage in a manner similar to other MutL proteins^58^. Thus, it is possible that MutLβ is recruited to the expanded CAG repeat DNA locus by MutSβ^59^, and such recruitment is necessary for the subsequent repeat expansion process (Fig. 8F). A recent study revealed a novel function of the mammalian MutSβ-MutLβ complex: it selectively binds to RNA:DNA hybrids (R-Loops) stabilized by G-quadraplexes, which then recruit the helicase FANCJ to restart the stalled DNA replication fork^60^. Thus, it is possible that a similar mechanism operates on the expanded CAG repeats that are prone to R-loop formation^61^. Our study does not exclude a potential role for Pms2 in mHtt CAG repeat expansion and pathogenesis, but genetically it is not necessary for such a process as it may be functionally redundant with Mlh3. Prior studies have established an essential role for Mlh3 in driving mHtt CAG expansion in Q111 KI mice^62,63^. Putting it together, our study suggests a revision of the previous mechanistic model for HD GWAS/MMR genes in driving somatic CAG expansion and downstream pathogenesis, in which the endonuclease negative MMR complex (MutSβ-MutLβ) and at one or both endonuclease positive MMR complexes (MutSβ-MutLγ and/or MutSβ-MutLα) would be needed for the process (Fig. 8F).

### Sequential striatal and cortical pathogenesis in HD mice requires Msh3

An important unresolved question in the HD field is to what extent the genetic manipulation of HD GWAS/MMR genes such as Msh3 could benefit the disease, and if so, could long-term Msh3-reduction exert sustained benefit without inducing harmful effects in HD. Currently, Msh3 genetic reduction in HD mice stabilizes somatic mHtt CAG repeat expansion and/or reduces nuclear mHtt accumulation in the striatal neurons^29,64^. It remains unclear whether any cortical HD mouse phenotypes depend on Msh3. In HD postmortem brains, although a subset of striatal and cortical pyramidal neuronal types show CAG repeat expansion and degeneration, MutSβ is selectively expressed at higher levels in the striatal MSNs^39^ but not cortical neurons^42^, suggesting a potential rate limiting role of Msh3 expression levels in conferring neuronal vulnerability in HD. In this study, we first performed deep phenotyping of Q140 KI mice (expressing endogenous CAG-expanded mHtt) over a long age range to define the sequential steps in pathogenesis from the early onset and progression of striatal pathogenic phenotypes to the later onset and progression of cortical phenotypes. The brain regional and cell-type selectivity and temporal progression of these pathological phenotypes in this HD mouse model mimic the selective disease patterns in HD^1^. Importantly, we show that the pathogenic triad (CAG instability, mHtt aggregation, and transcriptionopathy) follows a temporal order of onset, i.e., striatal MSNs at 6m, piriform cortical neurons at 12m, and motor-sensory cortical neurons at >12m (i.e., 16m-20m). Importantly, these molecular and cellular pathological phenotypes are associated with other neuropathological deficits including synaptic protein (Actn2) loss at 12m, upregulation of GFAP+ astrocytosis at 20m, and locomotor deficits. Thus, our study suggests that the expression of CAG-expanded endogenous mHtt can elicit lifelong disease course that has brain regional and neuronal cell type specificity, and overall disease progressivity, that is partially recapitulating those found in HD.

The second important conclusion from our study is that genetic reduction of Msh3 and Pms1 can prevent the onset of both striatal and cortical pathogenesis and overall behavioral deficits in Q140 mice. Since the onset of the striatal disease is at around 6m, and cortical disease is delayed by 6m or more (dependent on cortical regions), our study provides another important implication for potential therapeutics targeting the Msh3 and other MMR genes: if the therapeutic intervention is too late to counteract the striatal disease due to long CAG expansion, there can still be benefits in brain regions with a slower rate of repeat expansion and later age of disease onset. Thus, MMR targeting therapeutics could be potentially helpful in preventing disease onset or slowing disease progression, especially when such disease processes are attributed to neuronal cell types with different rates of repeat expansion.

We do recognize some of the limitations of the HD mouse models. They do not recapitulate the full scale of disease (e.g. frank neuronal loss and movement disorders) found in HD patients, and the inherited CAG repeat length unusually need to be much longer (e.g. >100 CAG) than those found in the patients to observe the neuronal selective and progressive pathologies. The latter is also true for other CAG repeat disorders such as SCA1^65,66^ and spinal bulbar muscular atrophy (SBMA)^67^, and may reflect a more tolerability of murine neurons to the repeat-induced toxicities or the relative short lifespan of the mouse models to fully manifest the disease.

Notwithstanding these limitations, our study demonstrates Q140 and possibly other HD mice with comparable CAG repeat length (e.g. BAC-CAG)^8^ could be valuable models to study the mHTT CAG-expansion driven progressive and selective pathogenesis that mimic aspects of selective neuronal vulnerability and disease progression in HD, and to evaluate how genetic or molecular perturbations could prevent or modify such longitudinal disease processes.

### Linear and high-rate of CAG-repeat migration is selective to MSNs in HD mice and depends on Msh3 gene-dosage

Prior studies have shown that striatal neurons in HD mouse models^68^ as well as MSNs and CPNs (but also a small subset of less vulnerable neurons) from HD patients have a higher propensity for somatic CAG repeat expansion^39,42^. However, the kinetics of such repeat expansion is thought to be exceedingly difficult to unravel due to the stochastic nature of CAG repeat expansion and highly expanded repeats being present only in a small subset of neurons^69–71^. In cultured patient iPSC-derived neurons, CAG expansion kinetics can be measured but the range of expansion (<6 repeats) and time of measurement are limited. Currently, there are no established approaches to quantify cell-type-specific somatic CAG expansion kinetics in various brain cells^72,73^.

In this context, a surprising and important finding in this study is that the purified MSN nuclei in Q140 show CAG expansion at a linear rate of +8.8 repeats per month. This result suggests that at a relatively high inherited CAG length, the repeat expansion is following a strictly linear, and highly synchronous rate with an adult onset (i.e. 2m) and over a 14m time-window. Moreover, Msh3 single or double null alleles also produce highly linear MSN modal CAG expansion at a reduced rate of 2.3 repeats/month and a highly retarded rate of 0.35 repeats/month, respectively. We deem this finding highly rigorous for multiple reasons: (i). the linear model fits the measured Mod-CAG in all three genotypes up to 16-20m of age; (ii). it quantitatively reports the Msh3-gene-dosage dependent slowing of CAG expansion in the MSNs; and (iii). the width of the CAG peak at a given repeat length is highly positively correlated with the modal CAG sizes, demonstrating the strict quantitative relationships between the MSN repeat expansion sizes and its distribution regardless of genotype and age. The latter also suggests the existence of very moderate asynchronous expansion during aging among the MSNs, but such effects are secondary to the overall synchronous repeat expansion. Moreover, we also showed that the inclusion of interneurons (FANS-NeuN) or total striatal tissue resulted in a CAG main peak at the inherited 140 CAGs. These results suggest the fast linear rate of CAG repeat migration at a cell-type population scale is selective to the MSNs, while striatal interneurons or other non-neuronal cell types still retain inherited CAG repeats in their cell populations. Future studies using FANS to purify nuclei from interneuron types or glial cell types will be needed to discern the CAG expansion kinetics in the other cell types in HD mice.

We are cognizant that our current study defines the first in vivo linear of mHtt CAG-repeat expansion in one of the vulnerable neuronal cell types in an HD mouse model, and there are many new questions that need to be addressed. First, our study opens the opportunity to determine modal CAG-repeat expansion rates in other neuronal and non-neuronal cell types in different brain regions in the Q140 mice, comprehensively testing the hypothesis that other brain cell types with different rates of repeat migration may predict the onset of pathologies (or lack thereof) in these cells. Second, additional studies can be readily pursued to examine whether different inherited CAG repeat lengths (i.e., allelic series mHtt KI mice) or human genomic mHTT transgene context (i.e., BAC-CAG)^8^ could confer different linear rates of CAG repeat migration in HD vulnerable neurons. Lastly, it would be important in the future to compare and contrast the findings from mouse models to those cell-type-specific repeat sizing in purified brain cell populations^39,42^ or single-cells^74^ from HD patients. The latter do not provide longitudinal kinetics of repeat expansion in various cell types, but modeling approaches^74^ and/or other repeat-length dependent molecular changes (e.g. transcriptome, mHtt aggregation) can be applied across model systems to strengthen the mechanistic understanding of somatic CAG expansion in HD vulnerable neurons.

### Evidence for somatic CAG-expansion threshold for mHtt aggregation in HD vulnerable neurons

A model in HD and the related repeat expansion disorder (RED) field is the two-stage disease process that posits first a somatic repeat expansion in vulnerable cell types with little cellular toxicities until the repeat length reaches a certain threshold that triggers the second phase with downstream toxicities and neuronal cell death^24^. A challenge to this model is a lack of direct experimental evidence to show a somatic CAG-repeat threshold-dependent toxicity phenotype in HD vulnerable neurons. A previous mathematical modeling of the relationships between inherited CAG genotype and age of HD onset and progression suggests a pathological threshold of mHTT in HD neurons is about 115 CAGs^69^. However, others have pointed out the wide 95% confidence interval of this threshold (70-165 CAGs) making the prediction difficult to interpret^72^. Thus, to causally define expanded CAG-length related pathological threshold, it would be essential to: (i) precisely measure neuronal cell-type-specific CAG repeat length and relate the quantitative data on neuropathology; and (ii) demonstrate that preventing CAG expansion beyond the threshold can prevent the onset of such pathology in a relevant neuronal cell type. To this end, our study provides the first definitive evidence that the formation of visible mHtt nuclear inclusions and neuropil aggregates, as detected by the commonly used anti-mHTT antibodies (i.e. PHP1 and EM48), requires CAG expansion beyond the inherited 140-CAG. Our conservative estimates of NI and NA thresholds are 150 CAGs and 192 CAGs, respectively. This result can be extended in the future to other neuronal cell types in Q140 and other KI mice, human genomic models of HD (BAC-CAG), and importantly, to HD patient post-mortem brain samples.

The thresholds defined in this study help explain prior findings in other HD mouse models. First, murine allelic series KI mice with CAG lengths ranging 92-185 show increasing propensity for striatal nuclear accumulation and aggregation, but only Q140 and Q175 (with 185 CAGs) mice show predominant NIs by 6m of age, consistent with their CAG repeat sizes just below or above the first NI-Threshold. Second, BAC-CAG mice with longest CAG repeats (120 CAGs) shows the earliest, striatum-selective NI formation^8^, while those human mHTT transgenic models with stable 97 or 128 CAA-CAG repeats do not show NIs by 12m of age^46,75^. Moreover, BAC model with a stable 226 CAA-CAG repeats does show prevalent nuclear inclusions in the striatum, suggesting mHTT with a long polyglutamine protein stretch, beyond that encoded by our CAG expansion thresholds (i.e., 150 and 190 repeats), is sufficient to elicit nuclear and cytoplasmic aggregation pathologies and severe disease-like behavioral and pathological phenotypes^45^. Moreover, consistent with our aggregation threshold findings, Q175 mice crossed to Msh3-hom show stabilization of CAG repeats but not the prevention of mHtt aggregation or transcriptional changes (Aldous *et al.*, 2024). This result is consistent with the model that the inherited 185 CAG repeats in Q175 already exceed the aggregation threshold of 150 and hence the stabilization of CAG repeats does not prevent mHtt aggregation.

### A causal relationship between somatic CAG-expansion and transcriptionopathy in HD mice

Transcriptionopathy is a common molecular pathological feature shared between HD mouse models and patients^76^, and genes dysregulated in HD mouse models and patients partially overlap^35,39,77^. In allelic series mHtt KI mice with increasing inherited CAG repeat length, transcriptionopathy caused by germline CAG repeat expansion is strongest in the striatum compared to other brain regions^6^. Cell-type-specific transcriptomic analyses reveal that the striatal MSNs are the most dysregulated in HD mice^35^ (Fig. 2) and HD patients^39^. Mechanistically, mHTT is known to interact with other transcription and chromatin factors (e.g., SP1, REST, CBP, FoxP2), especially through its glutamine-rich domains, to elicit transcriptional changes^78–82^. However, to date, there is no genetic perturbation evidence that targeting these nuclear transcription or chromatin factors can robustly influence the striatal MSN-selective, CAG-length dependent transcriptionopathy in HD mice.

Our current genetic screen, using striatal bulk tissue RNA-seq as a primary readout, enables us to rigorously test the roles of a majority of HD GWAS/MMR genes in mHtt-induced transcriptionopathy in vivo. Our results show that Msh3 and Pms1 are the main genetic drivers of such molecular pathology, with Msh2 and Mlh1 also playing a modifier role. This conclusion is supported by (i). a robust inverse relationship between CAG repeat instability and transcriptomic reversal in all our KO perturbations in Q140 mice; and (ii). our finding that the Q140/Msh3-hom with stable 140 CAG repeats only elicits transcriptional deficits that are substantially weaker and have a later onset than those in Q140 with expanding CAG repeats. Thus, at 140 CAG repeats, transcriptionopathy is mostly but not entirely dependent on CAG expansion. This study shows that transcriptionopathy does not appear to depend on an absolute threshold that is above 140-CAG, but our study cannot exclude a threshold below 140 CAGs exists and is needed for transcriptional deficits in MSNs in HD mouse models.

A recent preprint study^74^ of human postmortem brain tissue using single-nuclei repeat sizing and RNA-seq revealed that single striatal MSNs with CAG repeats of 150-500 CAGs, but not those below 150 CAG, show robust transcriptional changes as well as evidence of neuronal loss^74^. However, another study shows FANS-purified MSNs with more moderate expansion (e.g. 60-100 CAGs) still show elicit marked transcriptional changes^39^. Future studies with different methodological approaches should be applied to both HD model systems and patient postmortem brains to more precisely discern the relationships between MSN somatic CAG repeat sizes and the magnitude of transcriptionopathy as well as the exact genes that are dysregulated. Additionally, profiling chromatin dysregulation in vulnerable neurons from both HD mouse models and patients may further shed light on the mechanistic gap between CAG DNA repeat expansion and transcriptional dysregulation in vivo.

### Implications for therapeutic development for HD and related repeat expansion disorders

Our study provides important implications for developing novel therapeutics targeting HD GWAS genes and/or somatic CAG repeat instability in treating HD and related REDs.

First, our study suggests that therapeutic targeting in HD should focus on the genetic or molecular reduction of the levels of MSH3 and PMS1, functional inhibition of MutSβ-MutLβ interaction, or inhibition of the latter complex’s binding to the disease-relevant DNA substrates.

Second, our study provides an integrated preclinical platform, including rich molecular, pathological, and behavioral readouts, to demonstrate the long-term in vivo efficacy as well as safety of targeting key MMR genes such as Msh3 and Pms1 for disease mitigation. Our study demonstrates the benefit of targeting Msh3 extends beyond striatal neurons and could help prevent late onset pathogenesis in other brain regions such as the cortex, which was not known prior to this study. The highly quantitative linear relationship between modal CAG repeats in MSNs and age in Q140 MSNs, and its exquisite sensitivity to Msh3 and Pms1 gene reductions, provide robust benchmarks to test future genetic and molecular therapeutics against these prime HD GWAS targets that are known to have relatively low cancer liability^21^. Since Q140 has a CAG repeat size just below the threshold for mHtt aggregation and its transcriptionopathy is predominantly driven by MMR genes, we propose that Q140 or other HD mouse models with comparable inherited CAG repeat lengths (e.g. BAC-CAG) could be optimal and robust models to test intervention to stabilize or reduce CAG repeats via MMR-targeting or other molecular tools^83^.

Our study provides candidate biomarkers that could be further explored in mouse models and patients. We showed that mHtt aggregation in MSNs is driven by CAG expansion thresholds, i.e., about 150 CAGs. If validated in the patients, PET ligand for mHTT aggregates could be used as a surrogate biomarker for somatic CAG expansion^84^, at least in early stages of the disease. Our study also defined 479 genes that are highly associated with MSN modal CAG repeat length in striatal MSNs, regardless of ages or genotypes of the tissues. It is interesting that multiple Mod-CAG associated genes, e.g. upregulation of Onecut1 and downregulation of Pde10a and Ano3, are also found among significant DEGs in the same direction in HD MSNs^39^. Future studies will be needed to further define the genes that are commonly dysregulated in both HD mouse models and patients^39,74^, and asses their dependency on somatic CAG-repeat length. Such commonly dysregulated genes could provide candidate biomarkers for somatic CAG expansion in HD vulnerable neurons. It is worth noting that our Mod-CAG dependent genes include well known clinical PET ligands that are shown to be altered in HD patients, i.e. PDE10A, DRD1, DRD2, OPRK1^54^. Other candidate biomarker proteins include secreted or transmembrane proteins that could be found in the CSF (e.g., FST, IGFBP4). Mining mHTT somatic expansion/MMR-gene-dependent transcriptomes and proteomes in HD mouse models could provide a rich source for biomarkers for MMR-targeting therapies for HD.

Lastly, there are more than 40 human repeat expansion disorders (REDs) caused by short tandem repeat instability^85,86^, including but not limited to other CAG-repeat disorders, such as myotonic dystrophy, fragile X, Friedreich Ataxia etc. Moreover, DNA repair genes such as *MSH3* or *FAN1* have been shown to be within modifying GWAS loci for several other REDs including X-linked Dystonia-Parkinsonism (XDP)^87^, Myotonic Dystrophy type 1 (Flower *et al*, 2019), and some spinocerebellar ataxias^88^. Thus, the genetic modeling, mechanistic findings, and therapeutic enablement stemmed from this study could help advance the mechanisms and therapies for other REDs as well.

## ACKNOWLEDGMENTS

We thank the UCLA Neuroscience Genomics Core (UNGC) and UCLA Technology Center for Genomics & Bioinformatics (TCGB) for RNA-seq data acquisition; the UCLA Flow Cytometry Core for the FACS service; the Jackson Laboratories, KOMP/MMRRC at UC Davis, the National Cancer Institute Mouse Repository, and Rodent Genetics Core at Cedar-Sinai for generating new CRISPR/Cas9 KO mice, supplying and rederiving of existing KO mice. The research described in this study was supported by CHDI Foundation, Inc. Huntington’s Disease (HD) research in the Yang lab is also supported by NINDS/NIH grants (R01NS113612), Hereditary Disease Foundation, and donations from the HD patient families. X.W.Y is supported by the Terry Semel Chair in Alzheimer’s Research and Treatment at UCLA.

## AUTHORS CONTRIBUTIONS

X.W.Y. conceptualized and directed the entire study. X.W.Y., N.W., and P.L. wrote the manuscript. X.W.Y., N.W., S.Z., and P.L. designed the detailed experiments and data analysis pipeline, and performed data analyses and interpretation. J.R., J.A., B.P., T.V., and S.H. provided inputs into the experimental design and data analysis/interpretation. N.W. (with help from S.Z.) led a group of researchers in the Yang lab, including L.R., M.P., R.V., X.G., Li.D. and Le.D., to import or generated KO mouse lines, bred the KO mice with Q140 KI mice, performed mouse colony management, dissected mouse tissues, prepared bulk RNA or single nuclei-RNA for RNA-seq, prepared genomic DNA for repeat instability assays, conducted neuropathological and behavioral studies. F.G. helped perform RNA-seq data processing. P.L. performed all the RNA-seq related data analysis and generated the plots and tables for the manuscript. N.W., S.Z, L.R. and Le.D. performed the data analyses for repeat instability, neuropathology, and behavioral tests.

## DECALRATION OF INTERESTS

X.W.Y. is a Scientific Advisory Board (SAB) member for Lyterian Therapeutics and Ophidion Therapeutics, and a former SAB member for Mitokinin Therapeutics and Triplet Therapeutics. X.W.Y. has served as a consultant for Ionis, Biogen, Novartis, Roche, LifeEdit Therapeutics, PTC Therapeutics, Ascidian Therapeutics, Sangamo Therapeutics, and Forbion. P.L. served as occasional bioinformatics consultant for The Bioinformatics CRO, Inc; Vynance Technologies, LLC; and FOXO Technologies, Inc.

## METHODS

| REAGENT or RESOURCE | SOURCE | IDENTIFIER |
| --- | --- | --- |
| **<u>Experimental Models: Organisms/Strains</u>** |  |  |
| Mouse: Msh3 KO | NCI | B6.129-*Msh3*<sup>tm1Rak</sup>/Nci |
| Mouse: Mlh1 KO | NCI | B6.129-*Mlh1*<sup>tm1Rak</sup>/Nci |
| Mouse: Msh6 KO | NCI | B6.129-*Msh6*<sup>tm1Rak</sup>/Nci |
| Mouse: Msh2 KO | JAX | RRID:IMSR_JAX:016231 |
| Mouse: Pms2 KO | JAX | RRID:IMSR_JAX:010945 |
| Mouse: Polq KO | JAX | RRID:IMSR_JAX:006194 |
| Mouse: Pms1 KO | MMRRC | RRID:MMRRC_049126-UCD |
| Mouse: Tcerg1 KO | MMRRC | RRID:MMRRC_050004-UCD |
| Mouse: Ccdc82 KO | Yang Lab (UCLA) | Available upon request |
| Mouse: Rgs9-Cre | JAX | RRID:IMSR_JAX:020550 |
| Mouse: Q140 | JAX | 400809 |

### Experimental animals

In this study, we bred KO mice for 9 genes (Table S1 and Key Resource Table) with HD KI Q140 mice (and Rgs9-Cre mice for the Msh2 cKO line) to generate respective heterozygous and homozygous (if available) KO mice in WT or Q140 heterozygous background. Briefly, 5 KO mouse lines were imported from public repositories (JAX or MMRRC/UCD); 3 KO lines were imported from NCI and cryo-recovered at Cedars Sinai; Ccdc82 KO was generated by Yang Lab using the CRISPR/Cas9 method at the Jackson Laboratory’s Customer Model Generation Core. For each crossing, offspring were weaned and tattooed with ear tags at around 3 weeks of age, and genomic DNAs extracted from ear biopsies were used for genotyping PCR. All mice were maintained and bred under standard conditions consistent with National Institutes of Health guidelines and approved by the University of California, Los Angeles Institutional Animal Care and Use Committees. The cages were maintained on a 12:12 light/dark cycle, with food and water ad lib. Locomotor behavior was measured in all mice using open-field test as previously described one week before euthanizing for experiments.

## METHOD DETAILS

### Tissue collection and sample preparation

Mice were deeply anesthetized with Nembutal (50 mg/kg IP). For RNA-seq and western blot assays, anesthetized mice were transcardially perfused with DEPC treated 1xPBS, the cortex, striatum, cerebellum and major peripheral organs were then separated, snap frozen on dry ice and stored at −80°C until further processing. For immunohistochemistry staining, anesthetized mice were transcardially perfused with 1xPBS, followed by 4% paraformaldehyde (PFA) in PBS, pH 7.4. The brains were dissected and post-fixed with 4% PFA overnight at 4 °C. Subsequently, the brains were transferred to tubes containing 30% sucrose in cold 0.1M PBS. After they sank to the bottom of the tubes, the brains were removed, briefly drained and snap frozen on dry ice cold 2-methylbutane.

### Immunohistochemical staining and imaging

Coronal brain sections were cut at 30 µm with Leica Cryostat and stored at −20°C in cryopreservation solution (30% sucrose, 30% glycerol in 1xPBS) for further processing. Immunostaining was carried out using published methods (Gu et al., 2022). PHP1 antibody (Millipore MABN2490; 1:2,000 dilution) and EM48 (Millipore MAB5374; 1:350 dilution) was used for mHTT aggregate detection; Darpp32 antibody (Cell Signaling 2306; 1:500 dilution) was used for detection of MSNs; Gfap antibody (Dako Z0334; 1:1000 dilution) was used to detect active astrocytes; Actn2 antibody (Abcam ab68167, 1:500) was used to quantify alpha-actinin 2 expression in striatum. Images were acquired using a Dragonfly High-Speed Confocal Microscope 200 and the Fusion 2.0 software package (Andor, Oxford Instrument). Laser and detector settings were maintained at constant for the acquisition of each immunostaining. For all analyses, at least three mice per genotype and 3 images per brain using were taken with 60x oil objective lens, at 2,048 × 2,048-pixel resolution, with z-step size of 2.2 μm at 30 μm thickness.

### Analysis of CAG repeat instability in bulk tissues

About 100 µg tissues of cortex, striatum, cerebellum, liver and tail, dissected from 5-6 mice per group were digested with tissue digestion buffer containing proteinase K overnight in a 55°C water bath. Genomic DNA was isolated using phenol: chloroform method. CAG instability assay was performed at Laragen Inc. Briefly, the forward primer was fluorescently labeled with 6-FAM (Applied Biosystems) and products were resolved using the ABI 3730xl DNA analyzer (Applied Biosystems) with GeneScan 500 LIZ as internal size standard (Applied Biosystems). GeneMapper v5 (Applied Biosystems) was used to generate CAG repeat size distribution traces. Expansion and contraction indices were quantified from the GeneMapper CAG repeat distributions as reported (Lee, et al. 2010). Briefly, the highest peak in each trace was used to determine a relative threshold of 10% and peaks falling below this threshold were excluded from analysis. Peak heights normalized to the sum of all peak heights were multiplied by the change in CAG length of each peak, relative to the highest peak, and summed to generate an instability index representing the mean repeat length change.

### Analysis of CAG repeat instability in striatal Ctip2+ nuclei

Frozen mouse striatal tissues were transferred to a 1.5-ml microcentrifuge tube, with 300 μl NP40 Lysis Buffer (10mM Tris-HCl pH7.4, 10mM NaCl, 3mM MgCl2, 0.1% Nonidet P40 Substitute, 1mM DTT in nuclease-free water) added and homogenized 15x using a Pellet Pestle on ice. Another 1 ml NP40 Lysis Buffer was added, and samples were incubated for 5 min on ice with pipette mixed twice during incubation. The lysates were passed through a 70 μm strainer into a 2-ml tube and centrifuged at 500 rcf for 5 min at 4°C. Most of the supernatant was removed, then 1 ml PBS w/ 1% BSA was generally added to the pellet without mixing. After incubation for 5 min on ice, the pellet was resuspended by pipette mixing. With another centrifuge at 500 rcf for 5 min at 4°C, most supernatant was removed, and the pellet was resuspended with 1ml PBS w/ 1% BSA and 1% Normal Goat Serum. 1:500 anti-Ctip2-FTIC (Abcam ab123449) was added to all the sample tubes except the negative control. Samples were rotated 45 minutes at 4°C, then filtered into FACS tubes and collected by Fluorescence-activated nuclei sorting (FANS) with 70um nozzle. Genomic DNAs were extracted from sorted nuclei using PureLink Genomic DNA Mini Kit (Invitrogen) following manufacture protocol. CAG instability assay was performed at Laragen Inc. as described above.

CAG peak location (Mod-CAG) was defined as the CAG length at the highest value of the CAG PCR trace at or above the tail CAG length. A baseline value was defined as the minimum of the trace and the peak height relative to baseline was calculated as the difference of the absolute peak height and the baseline level. We chose a fraction of the relative peak height (say 0.1 or 10%) and defined the corresponding fraction height (e.g., 10% of the peak height relative to baseline level). Starting from the peak location, the left and right peak boundaries were defined as the points in which, of 10 consecutive trace (height) values, at least 5 were below the fraction height. The reason for considering 10 consecutive trace values is to smooth out local fluctuations. The peak width was then defined as the difference of CAG values at the right and left peak boundaries.

### Quantitation of Htt expression by western blots

To test whether deleting DNA repair genes change Htt protein levels, these proteins were quantified by western blot assay, probed with anti-Htt (Sigma-Aldrich MAB2166, 1:2,000 dilution) and α-tubulin (Proteintech 66031-1, 1:10,000). Briefly, brains from 3-month old mice, of each line, were dissected, 40 µg protein from lysates of striatum from each brain was mixed with NuPAGE LDS loading buffer (Invitrogen), and heated for 10 min at 70°C, then resolved on 3-8% Tris-Acetate NuPAGE gel (Invitrogen). After protein transfer to PVDF membrane, blots were probed with the respective antibodies. Densitometric values from scanned Western blot films were obtained using ImageQuantTL software (GE Healthcare).

### Open field test for locomotion behavior

Mice were individually video-recorded under deem light in a clear plexiglass behavioral recording chamber (40 cm x 40 cm x 40 cm) with a transparent floor through which an angled mirror allowed video-recording of behavioral activity from below. This bottom-up view allowed for clear, unobstructed observation and analysis of even the most subtle or slightest of mouse behaviors. Mice were video-recorded for 15 minutes, and DeepLabCut Toolbox (https://github.com/DeepLabCut/DeepLabCut) was used for behavioral quantification.

### RNA extraction and bulk RNA-seq

Frozen tissues were lysed in Trizol (Invitrogen; Carlsbad, CA) and total RNA was extracted using Qiagen (Valencia, CA) RNeasy kit with QIAshredder columns, per manufacturer recommendations, including on-column DNase digestion. cDNA libraries were generated using Illumina TruSeq RNA Library Prep Kit v2 and sequenced on Illumina HiSeq 4000 or NoveSeq S4 sequencer with a minimum read depth of 37.5 million/sample. Clipped reads were aligned to mouse genome mm10 using the STAR aligner using default settings. Read counts for individual genes were obtained using HTSeq. RNAseq data has been deposited to the Gene Expression Omnibus (GEO) repository (www.ncbi.nlm.nih.gov/geo).

### Bulk RNA-seq data preprocessing

Each cross was analyzed as a separate data set. In each data set, we retained only genes with at least 0.5 robust count per million reads (CPM) in at least the number of samples in the smallest group (genotype with smallest number of samples, generally 8). The rationale for is to only include genes that are likely to be expressed in at least one genotype. The robust CPM is calculated by first normalizing the counts using the Relative Log Expression (RLE) normalization implemented in DESeq2 and then calculating “counts per million reads” from the normalized data. To identify potential outliers, we used a modified version of the sample network methodology originally described in ^1^. Specifically, to quantify inter-sample connectivity, we first transformed the raw counts using variance stabilization (R function varianceStabilizingTransformation which also includes RLE normalization) and then used Euclidean inter-sample distance based on the scaled profiles of the 8000 genes with highest mean expression. The intersample connectivities k were transformed to Z scores using robust standardization,

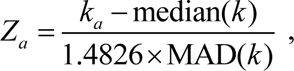

where index a labels samples, MAD is the median absolute deviation, a robust analog of standard deviation, and the constant 1.4826 ensures asymptotic consistency (approximate equality of MAD and standard deviation for large, normally distributed samples). Finally, samples with Z_a_ < −5 were removed.

### DE testing

To make DE testing robust against potential outlier measurements (counts) that may remain even after outlier sample removal, we calculated individual observation weights designed to downweigh potential outliers. The weights are constructed separately for each gene. First, Tukey bi-square-like weights *λ* ^2^ are calculated for each (variance-stabilized) observation *x_a_* (index *a* labels samples) as

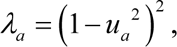

where

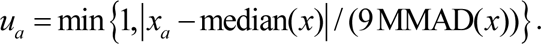

The median is calculated separately for each gene across all samples. MMAD stands for modified MAD, calculated as follows. For each gene, we first set MMAD = MAD. The following conditions are then checked separately for each gene: (1) 10^th^ percentile of the weights λ is at least 0.1 (that is, the proportion of observations with weights <0.1 is less than 10%) ^3^ and (2) for each individual genotype, 40^th^ percentile of the weights λ is at least 0.9 (that is, at least 40% of the observation have a weight ≥ 0.9). If both conditions are met, MMAD = MAD. If either condition is not met, MMAD equals the lowest value for which both conditions are met. The rationale is to exclude outliers but ensure that the number of outliers is not too large either overall or in each genotype group. This approach has previously been used in ^4,5^.

In a preliminary analysis of the Q140/Rgs9-Cre/Msh2 fl-hom and -het cross we found a relatively large overall variation component unrelated to genotype (or sex). Hence, we used the Surrogate Variable Analysis (SVA, R package sva ^6^) to calculate latent factors known as Surrogate Variables (SVs) representing unwanted variation and used the first SV as a covariate in DE testing.

DE analysis was carried out using DESeq2 (Love et al., 2014) version 1.36.0 with default arguments except for disabling outlier replacement (since we use weights to downweigh potential outliers) and independent filtering (since we have pre-filtered genes based on expression levels). For 12-month Rgs9-Cre/Msh2 fl-het and -homo data we used first SV as a covariate. Genes DE at FDR<0.1 were called significant.

### Measures of overall transcriptome-wide reversal and exacerbation

We use several statistics to quantify the extent of transcriptomic reversal for each perturbation. They can be broadly divided into two groups, statistics based on significance and statistics based on fold change. For the former, for each perturbation (KO) we report the numbers of significantly reversed or exacerbated genes at a defined significance threshold (e.g., FDR<0.1), defined as genes that are significantly DE in a baseline Q140 vs. WT test as well as in Q140/KO vs. Q140. Of these, the reversed genes’ fold change has opposite signs in Q140 vs. WT and Q140/KO vs. Q140 (i.e., compared to Q140, expression in Q140/KO moved towards WT levels), while for exacerbated genes the fold changes have the same sign. Although numbers of significantly reversed or exacerbated genes are intuitive, they require selecting a significance threshold and at any given significance threshold they are strongly influenced by the numbers of samples and other technical factors. Another useful statistic is a transcriptome-wide correlation of Z statistics for Q140 vs. WT and Q140 vs. Q140/KO which we call the reversal correlation (of Z statistics). When this correlation is positive (negative), the KO perturbation causes expression changes in Q140 background that overall move expression levels closer to (further away from) WT levels. Although sample numbers affect the scale of Z statistics, the correlation is scale-independent and hence only sensitive to the number of samples in the sense that higher numbers of samples generally increase signal to noise ratio. One could also define an analogous reversal correlation for log fold changes. Using correlation as a measure of transcriptome-wide reversal is in many ways attractive but it by definition cannot measure the actual amount of expression reversal or exacerbation. Hence, we also defined a measure of overall log fold change normalization. Specifically, we define an “overall reversal fraction” as the coefficient of a weighted linear model regressing log fold changes in Q140 vs. Q140/KO on log fold changes in baseline Q140 vs. WT. The weights are constructed so as to give more weight to genes with stronger evidence of DE in baseline Q140 vs. WT. We chose the form

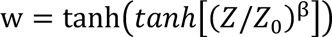

with *Z*_0_=3 and *β=*6. These settings result in weights that are low for *Z* ≤ 3, quickly rise for values around 3 and then flatten out, approaching 1 for *Z* much greater than 3. In our applications we find that overall rescue fractions are relatively insensitive to the choices of *Z*_0_ between 1 and 5 and *β* around 6. We further correct the regression coefficient for possible systematic differences between fold changes in Q140 vs. WT in the baseline data set and in the perturbation study that contains its own Q140 vs. WT test. Using the same weights, we calculate weighted variances of log fold changes for Q140 vs. WT in the baseline and in the perturbation data set and use their ratio to correct the overall rescue fraction. The rationale for this step is to account for potential dependence of estimated log fold changes on technical variables such as number of samples, biological and technical noise etc.

### Measures of individual gene reversal and exacerbation

For each individual gene, we define two statistics of reversal/exacerbation. First, the reversal score aims to identify genes with statistically significant DE for both Q140 vs. WT in baseline data and in Q140/KO vs. Q140. Denote the Z statistics for these two tests as *Z*_Q140-WT_ and *Z*_Q140/KO-Q140_. The reversal score is then defined as the minimum of |*Z*_Q140-WT_| and |*Z*_Q140/KO-Q140_|, multiplied by a sign that makes the reversal score positive when the two Z statistics have the opposite signs and negative otherwise. Thus, a reversal score of say 4 or more identifies genes that are DE at |*Z*| > 4 in both the baseline and the perturbation test and the DE is such that the perturbation reverses the effect of Q140 back towards WT levels. Since the reversal score is sensitive only to significance but not to the amount of reversal or exacerbation, we also define the reversal fraction as the fraction of the expression change between Q140 and WT that was reversed by the KO perturbation in Q140 background. Specifically, denoting the mean expression levels of a particular gene in WT, Q140 and Q140/KO genotypes by *E*_WT_, *E*_Q140_ and *E*_Q140/KO_, the reversal fraction is defined as *RF* = (*E*_Q140_-*E*_Q140/KO_)/(*E*_Q140_-*E*_WT_). This can also be expressed in terms of log_2_ fold changes (*lFC*) as

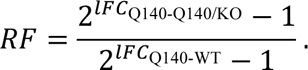

Positive reversal fraction between 0 and 1 means partial correction; reversal fraction is 1 for full correction and above 1 for over-correction. Negative values indicate exacerbation. Together, the reversal score and reversal fraction provide information both about statistical significance and the amount of expression correction. We note that the (individual gene) reversal fractions and (transcriptome-wide) overall reversal fraction, although clearly related, do not measure the reversal on the same scale: The individual gene reversal fraction is based on differences of mean expression values on the natural scale while overall reversal fraction is based on log fold changes, i.e., differences of log-transformed expression values.

Because all of the above reversal measures involve expression in Q140 genotype in two DE comparisons (Q140 vs. WT and Q140 vs. Q140/KO), using the same Q140 data in both comparisons results in a bias towards reversal in both transcriptome-wide and individual reversal statistics. To obtain unbiased reversal statistics, the two DE comparisons should be carried out on data from independent samples.

### Common analysis of Q140 vs. WT

We collected Q140 and WT samples from all of our bulk studies as well as from 52 perturbation studies at 6 months of age that gave rise to a large study of co-expression patterns in WT mouse striatum ^5^ and carried out 3 common Q140 vs. WT DE analyses: 810 samples (405 Q140 and 405 WT) samples from 52 crosses (referred to as GNV) at 6 months; 136 (66 Q140 and 70 WT) from the 9 independent 6-month Q140 and WT sets reported in this study, and 30 samples (17 Q140 and 13 WT) from 2 independent 12-month Q140 and WT sets also reported here. To account for potential batch effects, we tested DE for Q140 vs. WT in each data set separately (i.e., 52 sets in 6m GNV, 9 sets of 6m and 2 sets of 12m data in this study) and then combined the significance Z statistics for each gene using two approaches. First, we used Stouffer’s meta-analysis method ^7^ to obtain significance statistics that reflect the full number of samples used in each combined analysis. Second, we also combined Z statistics from individual studies using averaging which results in overall significance statistics “scaled” to (approximately) 8 samples per genotype and makes them comparable among the combined studies as well as to DE analyses in individual perturbations. To obtain overall log fold change for each gene, we averaged its log fold changes across all studies. In addition to overall meta-analysis and average, in 6m and 12m data from this study we also carried out meta-analyses and averages excluding each one of the data sets in turn. The aim of this “excluding one set” analysis is to obtain reference Q140 vs. WT statistics for each set (namely the excluded one) that are entirely independent from DE in Q140/KO vs. Q140 in that set.

### Enrichment analysis

Enrichment analyses were performed using our published methods ^5^. Briefly, enrichment calculations were carried out on sets of genes DE at FDR<0.1 or p<0.01 using our in-house R package anRichment (https://horvath.genetics.ucla.edu/html/CoexpressionNetwork/GeneAnnotation/) that implements standard Fisher exact test and a multiple-testing correction across all query and refence gene sets. To quantify the reversal/exacerbation effect of KO perturbations on Allelic Series WGCNA modules, we tested enrichment of genes DE in Q140/KO vs. Q140.

### Single-nucleus RNA-seq

Brain striatal tissues were dissected and snap-frozen on dry ice. Nuclei were isolated using the ‘Frankenstein’ protocol by Luciano Martelotto from University of Adelaide (DOI: dx.doi.org/10.17504/protocols.io.3fkgjkw). With Fluorescence-Activated Nuclei Sorting (FANS), 16,500 DAPI+ nuclei were collected for each sample and loaded onto the 10X chromium device. The library preparation was performed following 10X Genomics Chromium Single Cell 3’ Reagent Kits User Guide (v3.1). The pooled library was then sequenced with minimal 20,000 reads per cell on NovaSeq S4 platform at UCLA Technology Center for Genomics and Bioinformatics. Single nuclei (sn) -RNAseq data has been deposited to the Gene Expression Omnibus (GEO) repository (www.ncbi.nlm.nih.gov/geo), with the accession number **GSE241278**.

Starting from gene counts generated by CellRanger, we retained cells whose total expression (UMI count) is more than 1000 and in which there are at least 500 expressed genes (i.e., genes with a count > 0). We removed suspected doublets using DoubletFinder ^8^ with the doublet rate set to 0.08 based on information provided by 10x Genomics (https://kb.10xgenomics.com/hc/en-us/articles/360054599512-What-is-the-cell-multiplet-rate-when-using-the-3-CellPlex-Kit-for-Cell-Multiplexing-). The filtering and doublet removal were performed on cells from each mouse separately. We then combined data for the remaining cells across all samples into a single count matrix. We normalized the data using library size normalization, i.e., the normalized counts equal raw counts divided by the sum of UMI counts in the cell and multiplied by a scale factor that we chose to equal the median of per-cell UMI counts. For the initial clustering analysis we selected 4000 genes with highest adjusted variance ^9^ and added two categories of additional genes: marker genes for striatal cell types ^10,11^, microglial markers ^12^, combined MSN markers from sorted cell studies ^13–15^ with the restriction that their rank of decreasing adjusted variance is less than 10000, and 100 top hub genes from each of the strongly CAG length-dependent WGCNA gene modules from our study of an allelic series of mouse HD models with increased knock-in Htt CAG lengths ^16^, referred to as Allelic Series, with the restriction that their rank of decreasing adjusted variance is less than 15000. This procedure facilitates annotation of clusters found in subsequent analysis in cell type markers and Allelic Series WGCNA modules. We used singular value decomposition and projection to 60 leading principal components (PCs) to reduce noise in the data. Leuven clustering with resolution parameter = 2 was carried out on a nearest neighbor graphs constructed from 30 nearest neighbors of each cell. Clusters with at least 20 cells were labeled using positive integers, while cells in clusters with fewer than 20 cells were labeled by label 0 and considered not assigned to a cluster. We next tested differential expression between each cluster and the rest of the cells using Mann-Whitney-Wilcoxon rank sum test. We defined cluster markers as those genes that are overexpressed with p<10^−7^ and whose moderated log_2_ fold change is > log_2_(1.3). We define the moderated fold change as the difference between mean log_2p_ transformed expression within and outside of the cluster, where log_2p_(x) = log_2_(x+1). The +1 shift prevents large fold changes for low-expressed genes and hence ensures a certain minimum expression of cluster markers. For purposes of enrichment analysis, we limited the number of markers to the top 200 with strongest p-values. Conversely, if fewer than 20 genes passed these two thresholds, we dropped the fold change threshold and retained no more than 20 genes with strongest DE p-values that also pass p<10^−7^. We then carried out enrichment analysis of the cluster markers in cell type marker sets described in ^5^ to assign clusters to cell types. The initial clustering found 22 clusters, of which 17 exhibited strong enrichment (p<10^−8^) in at least of the cell type marker sets. For most of the 17 clusters, the enriched cell type was unique; only two relatively small clusters showed mixed enrichment (oligodendrocyte/microglia and oligodendrocyte/astrocyte). We next examined the maximum proportion of cells in each cluster that come from a single mouse and found that the 5 clusters without strong cell type annotation as well as the mixed oligo/microglia and oligo/astrocyte clusters and 1 other cluster (whose markers were enriched in astrocyte genes) had relatively high maximum proportion of cells from one mouse (> 0.12 while expected proportion is 1/24 ≈ 0.04). Hence, these clusters are more likely to reflect technical artifacts and we removed them from further analysis.

Our enrichment analysis was not sensitive enough to distinguish D1-from D2-MSN clusters. To distinguish them, we evaluated mean expression of D1 marker *Drd1* and D2 marker *Adora2a* in each cluster enriched in MSN marker genes (D1, D2 or general MSN). For each of the two marker genes, we ordered clusters by the mean expression, took the average (denote it *A*) of the second highest and second lowest cluster, and classified the expression as low or high depending on whether it was higher or lower than *A*. Clusters enriched in MSN markers that had a high expression of *Drd1* and low expression of *Adora2a* were labeled D1-MSN while MSN marker-enriched clusters with high expression of *Adora2a* and low expression of *Drd1* were labeled D2-MSN. In contrast, MSN marker-enriched clusters with either both high or both low expression of the two marker genes were labeled as general MSN. For D1- and D2-MSNs we found two clusters per cell type. For each MSN type, these two clusters were merged to form the final clustering with 12 clusters.

### Reversal analysis using nearest neighbors

For each cell *a* in any of the 6 genotypes we carried out the following procedure to find its 30 nearest neighbors among WT and Q140 cells. First, we excluded cells from the same mouse (i.e., if cell *a* is WT or Q140, the nearest neighbor candidates exclude cells from the same animal). Second, if there were more WT than Q140 candidates, we downsampled the WT cells to match the number of Q140 cells, and *vice-versa* if the number of Q140 candidates was larger. We then identified 30 nearest neighbors of cell *a* among the remaining candidates using Euclidean distance in the noise-reduced data reconstructed from the leading 60 PCs.

For each cell, we then calculated the proportion of WT cells in its 30 nearest WT and Q140 neighbors. For Q140 and WT genotypes, the difference in the proportion can be interpreted as a measure of effect size of the Q140 mutation. For other genotypes, the proportion can be considered an overall measure of location on an imaginary WT-Q140 line: values around 0 signify cells very similar to Q140 while values around 1 characterize cells similar to WT. To arrive at conservative significance estimates for differences, in each cluster we averaged the proportions of WT nearest neighbors by animal, resulting in 4 independent data points per genotype and cluster. To test for differences of the proportion among genotypes within each cluster, we used Tukey HSD test with *n*=4 per group.

### Pseudobulk DE analysis

To determine differentially expressed genes among the genotypes within each cluster, we created pseudobulk data for each cluster separately by summing the expression of each gene across cells from the same mouse and same cluster. We limited pseudobulking to those cluster/mouse combinations that had at least 30 cells.

We applied the bulk RNA-seq preprocessing and DE analysis as described above to the pseudobulk data sets, with the exception of requiring a robust CPM threshold that corresponds to approximately 10 UMI reads. Because of the widely varying numbers of cells and hence total gene expression in pseudobulk data, this translated to robust CPM thresholds ranging from ≈0.5 in D1-MSNs to over 100 in endothelial cells. We included he first Surrogate Variable calculated from variance-stabilized data as a covariate in DE testing.

In our analysis of genes commonly and individually reversed in D1-MSN and D2-MSN clusters, we simply defined commonly significantly reversed genes as those that were reversed at FDR<0.1 (i.e., DE at FDR<0.1 in both Q140 vs. WT and Q140 vs. Q140/Msh3-hom) in both C1_D1-MSN and C2_D2-MSN clusters. Genes significantly reversed in only one cluster were defined as those that pass the threshold in the cluster but not in the other, and genes that pass the threshold FDR<0.1 in neither of the clusters are called not significantly reversed in either cluster.

### Open-field locomotion test

The open field test is commonly used to measure locomotor activity. anxiety-like behavior and motor dysfunction and was performed similar as previous described ^5^. In our study the open field test was carried out 1h after lights off, nighttime open field (NTOF). Animals were transferred, in their home cage, in the dark, to the behavior room, which was illuminated under dim light. Animals were individually placed in the center of a square chamber and allowed to explore for 15 mins, with simultaneous video recording. The chambers were cleaned with 70% ethanol and dried between runs, to minimize olfactory cues. The video recordings were analyzed by DeepLabCut, an open-source Python package for quantifying animal behavior ^17^. Several behavioral parameters, including distance travelled, rearing time and rearing counts, time spent in the center and the periphery, as well as various gait measures (stance, splay and stride mismatch) were analyzed.

---

**Supplemental Figure S1.**
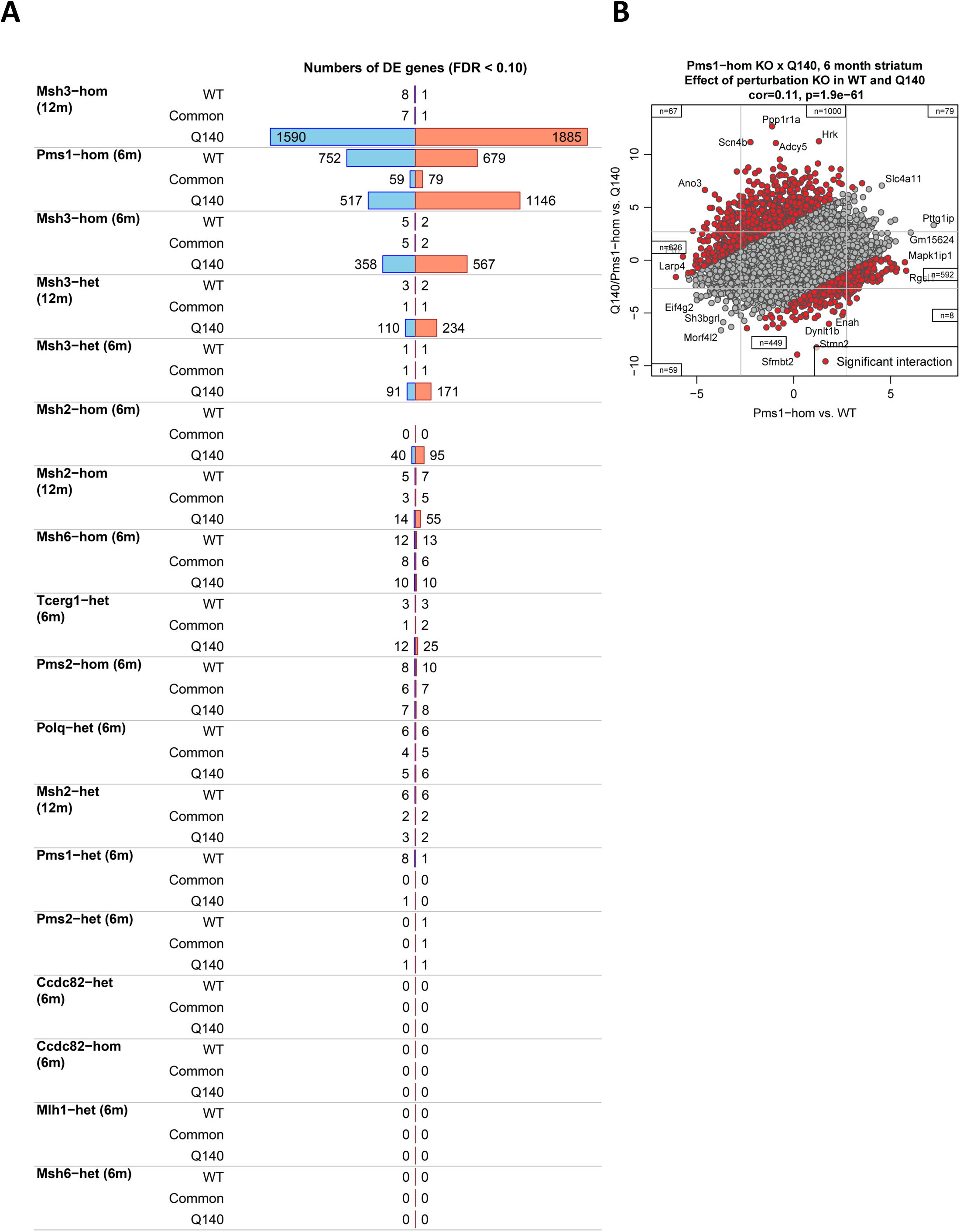
Striatal transcriptomic effects of HD GWAS/MMR KO alleles in the wildtype background. (A) Barplot representation of the number of genes significantly (FDR<0.1) DE in KO vs. WT (rows labeled WT), Q140/KO vs. Q140 (rows labeled Q140), and the overlap of the two for each perturbation. Blue and red bars represent down- and upregulated genes, respectively. Empty space in the WT row of Msh2-hom (6m) section indicates absence of data. (B) Scatterplot of DE Z statistics for Q140/Pms1-hom vs. Q140 (y-axis) against Pms1-hom vs. WT (x-axis). Each point represents a gene. Red color denotes genes with significant KO-Q140 interaction (i.e., significant KO-Q140 interaction term in the design of the negative binomial model regressing gene expression on genotype). Grey lines indicate approximate locations of FDR=0.1 thresholds. Numbers in the edge regions of the plot (these are regions with genes significant at FDR<0.1 in at least one of the two DE tests) give the number of genes.

**Supplemental Figure S2.**
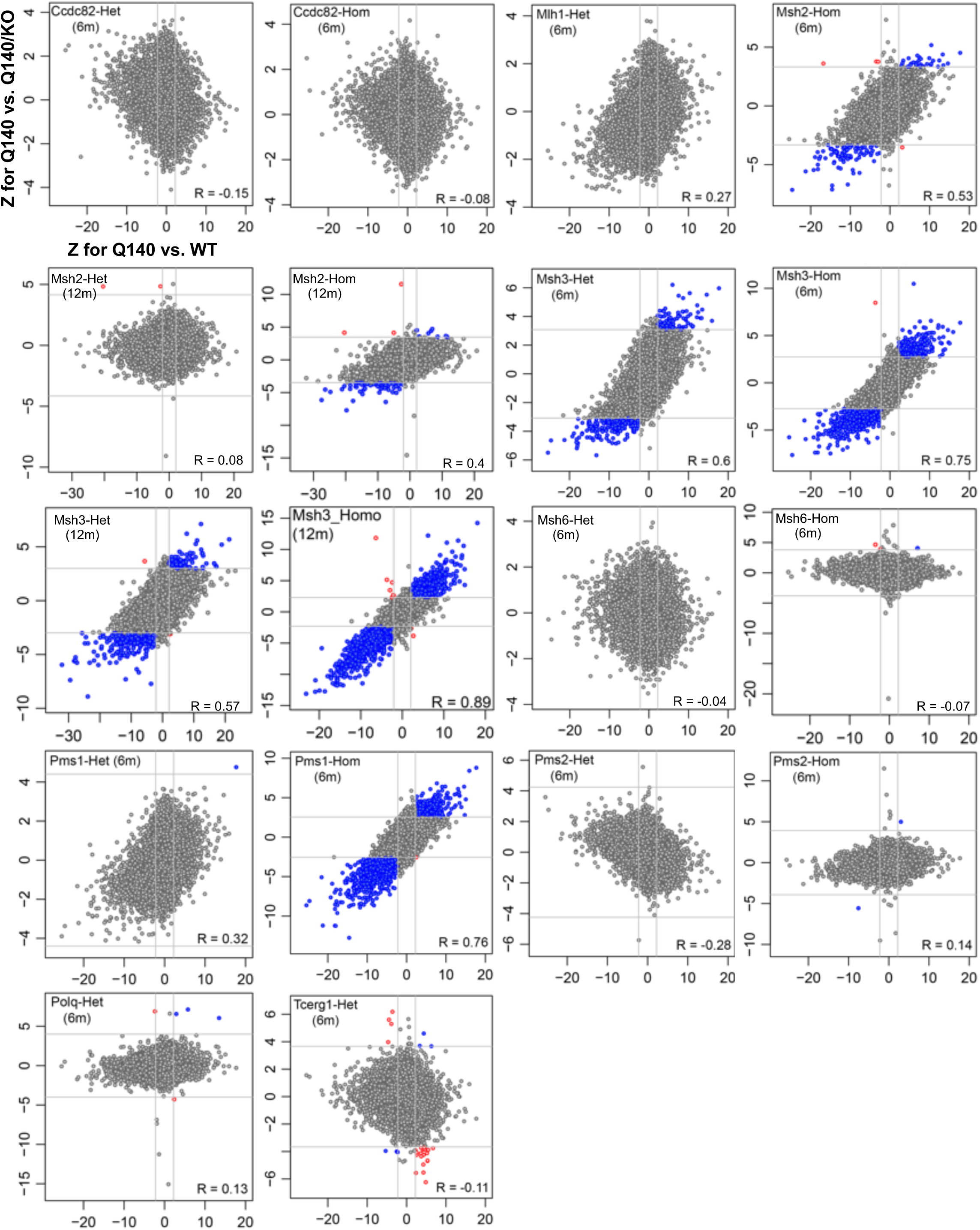
Scatterplots of Z statistics of transcriptomic reversal or exacerbation by the KO alleles in the Q140 background. Each panel shows a scatterplot of DE Z statistics for Q140 vs. Q140/KO for one perturbation (y-axis) against reference Q140 vs. WT (x-axis). Grey lines indicate approximate locations of FDR=0.1 thresholds for those tests where at least one gene passes the threshold. Blue and red color indicate significantly (FDR<0.1) reversed and exacerbated genes, respectively. Text within each plot indicates perturbation and the reversal correlation for the perturbation, i.e., correlation of x and y values in the plot.

**Supplemental Figure S3.**
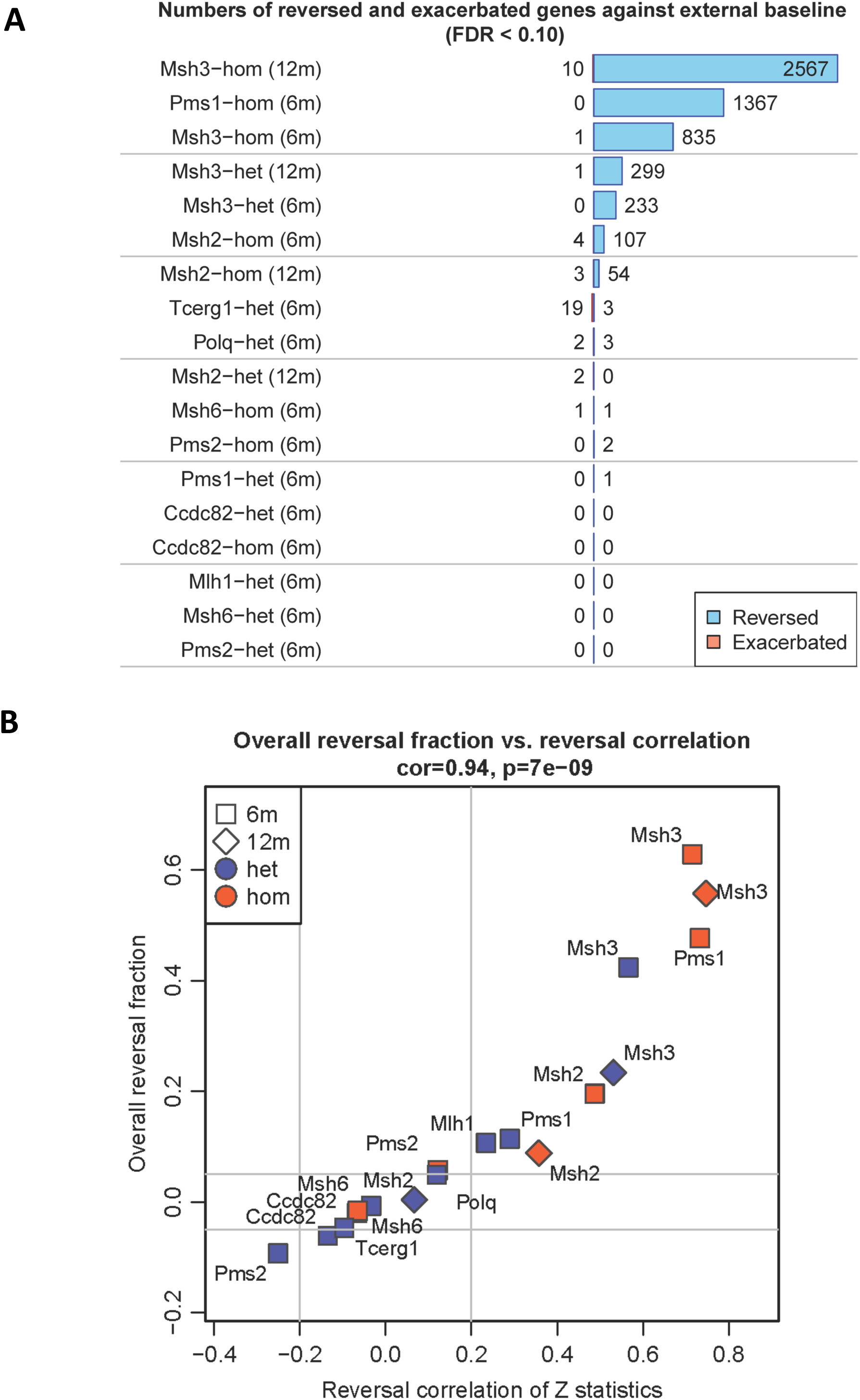
The number of reversed or exacerbated genes by Q140/KO perturbations and scatterplots of rescue fractions vs Z statistics. (A) Bars represent numbers of genes reversed (blue) and exacerbated (red) at FDR<0.1. (B) Scatterplot of overall rescue fraction (Methods, y-axis) vs. reversal correlation of Z statistics. Age and zygosity of each perturbation are indicated by point shape and color, respectively. Overall correlation and the corresponding significance (Student p-value) are indicated in the title of the panel.

**Supplemental Figure S4.**
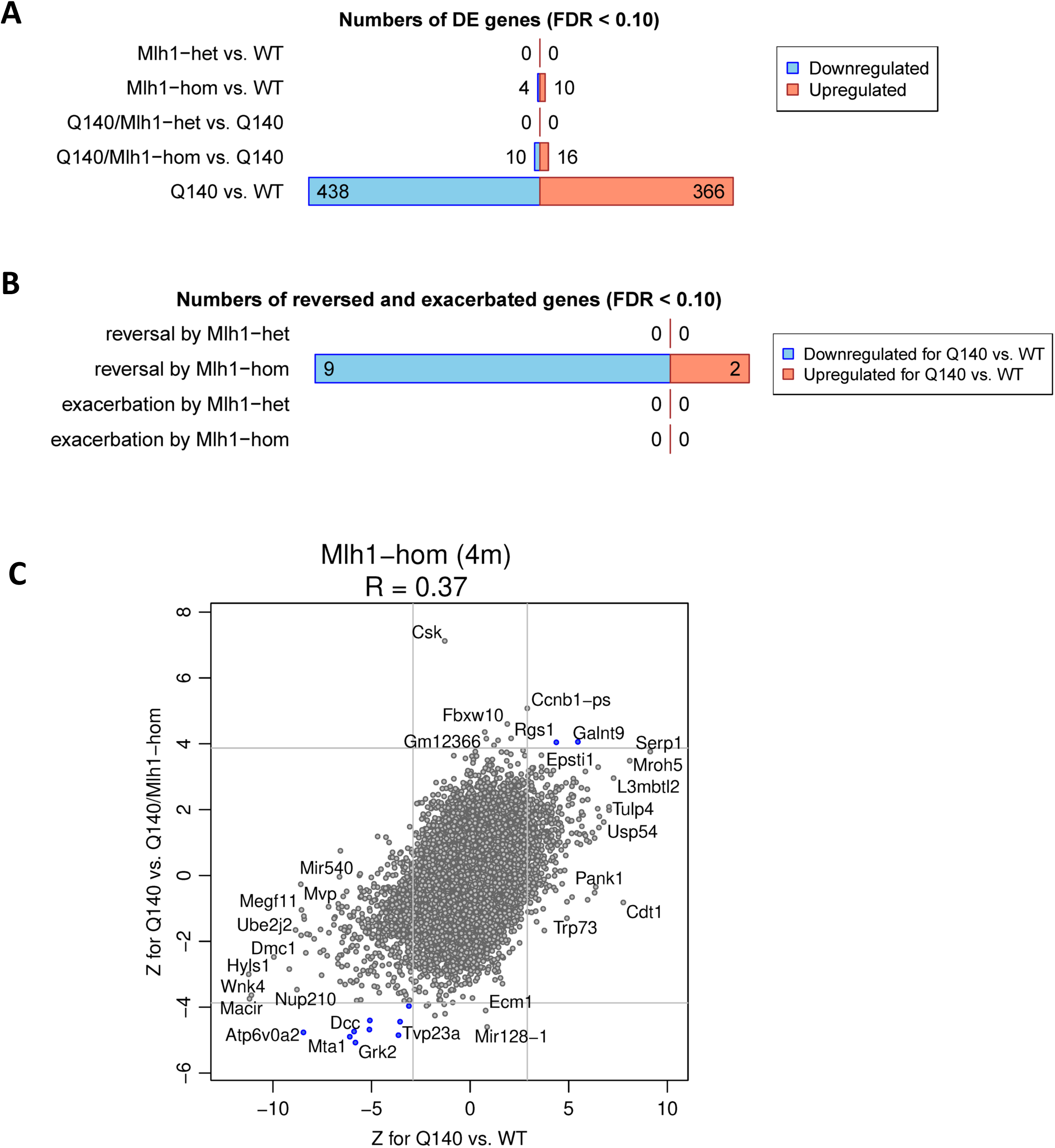
Mlh1 KO effects on Q140 striatal transcriptome at 4m of age. (A) Barplot of the numbers of genes significantly DE at FDR<0.1 in the tests indicated in row labels. Blue and red bars represent down- and upregulated genes, respectively. (B) Barplot of numbers of genes reversed and exacerbated at FDR<0.1. Blue and red bars represent genes down- and upregulated in Q140 vs. WT. (C) Reversal scatterplot for Mlh1-hom at 4m showing Z for Q140/Mlh1-hom vs. Q140 (y-axis) against reference Q140 vs. WT at 4m (x-axis). Grey lines indicate approximate locations of FDR=0.1 threshold. Blue points represent significantly reversed genes.

**Supplemental Figure S5.**
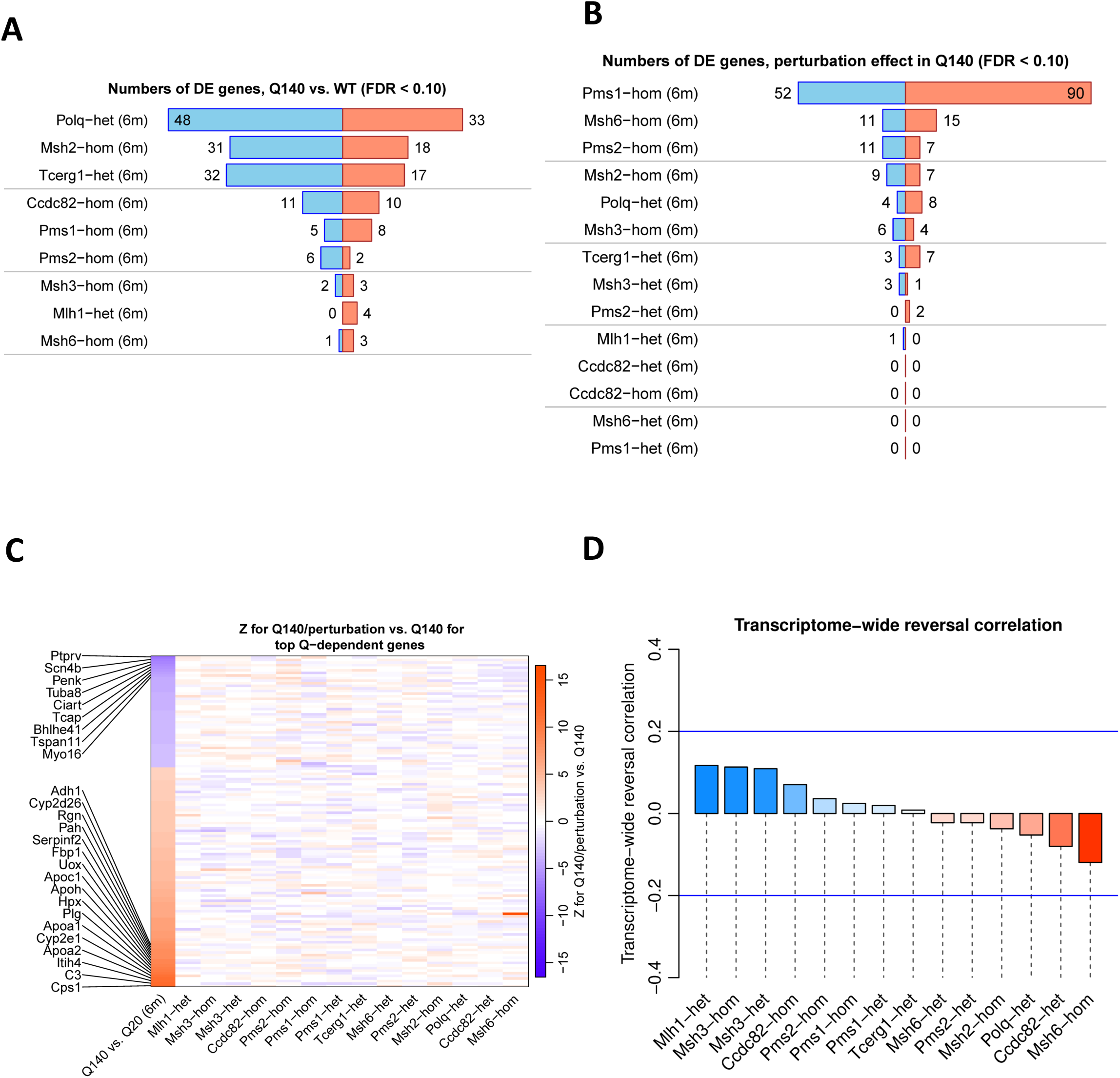
Cortex bulk RNA-seq for KO alleles in the Q140 background. (A) Numbers of significantly DE (FDR<0.1) genes for Q140 vs. WT at 6 in 9 independent data sets. Blue and red bars represent down- and upregulated genes, respectively. (B) Numbers of genes significantly reversed (blue bars) and exacerbated (red bars) by each perturbation at FDR<0.1. (C) Heatmap representation of DE Wald Z statistics for genes that pass FDR<0.1 and |FC|>1.2 in Q140 vs. Q20 at 6m in Allelic Series cortex. Columns include Q140 vs. Q20 in cortex at 6m in Allelic Series at Q140/KO vs. Q140 in perturbations at 6m. (D) Barplot of reversal Z correlations for cortex perturbations at 6m. Blue and red color indicate reversal (positive values) and exacerbation (negative values), respectively. Blue lines at ±0.2 indicate lower thresholds of what we consider moderate reversal/exacerbation. No 12m data included

**Supplemental Figure S6.**
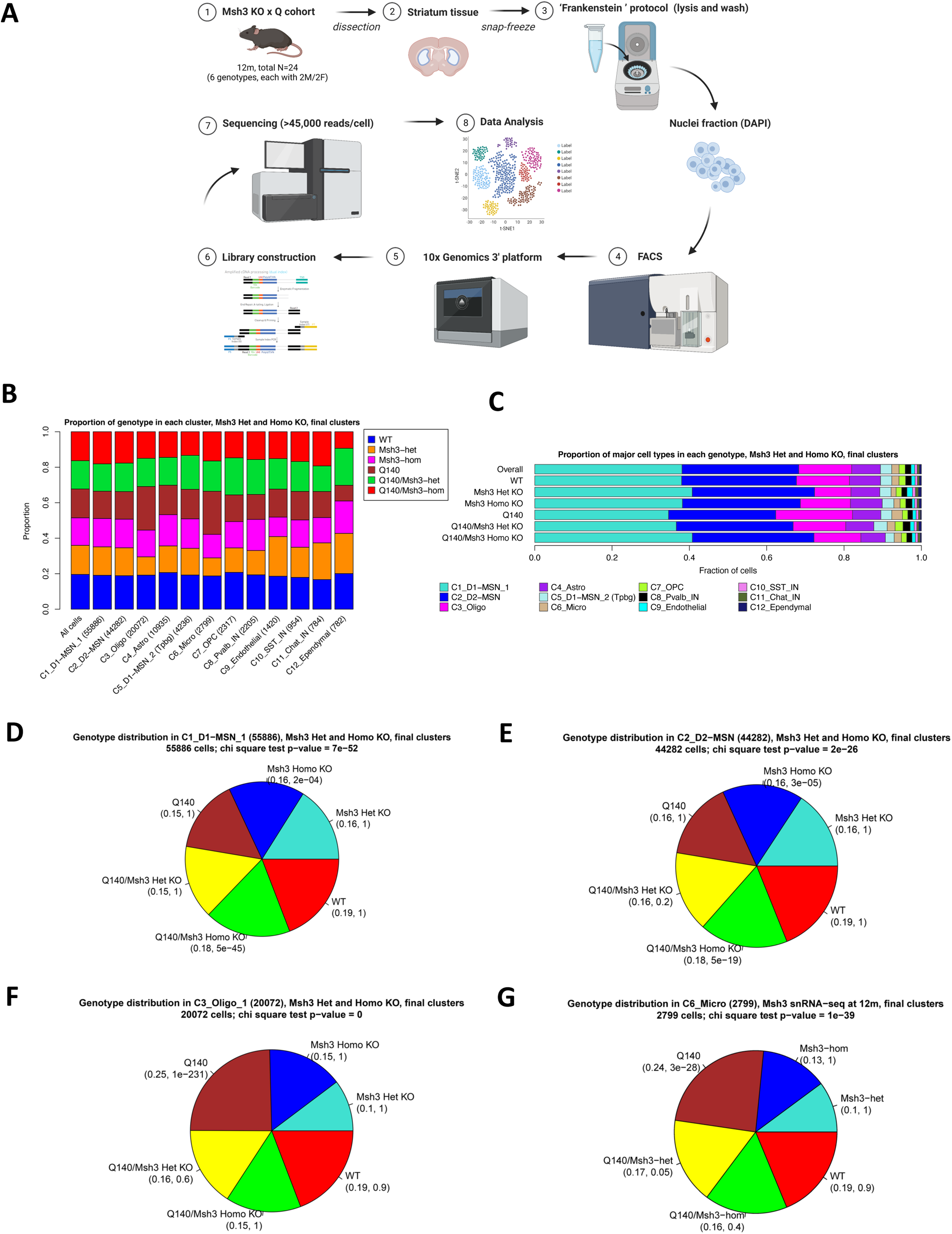
Single-nuclei RNA-seq of striatal nuclei from six genotypes in the Q140 cross to Msh3-KO at 12m of age. (A) Schematic of snRNA-seq pipeline. (B) Bars show proportions of the 6 genotypes overall (left-most bar) as well as in each cluster. (C) Bars represent the proportion of cells from each cluster in the 6 genotypes. (D) Proportions of genotypes among nuclei in C1_D1-MSN. Numbers in brackets below each genotype give the proportion and, in individual clusters, χ^2^ test p-value for over-representation of the genotype within the cluster (compared to the overall frequency of genotypes among all nuclei). (E) Proportions of genotypes among nuclei in C2_D2-MSN. (F) Proportions of genotypes among nuclei in C3_Oligo. (G) Proportions of genotypes among nuclei in C7_Micro.

**Supplemental Figure S7.**
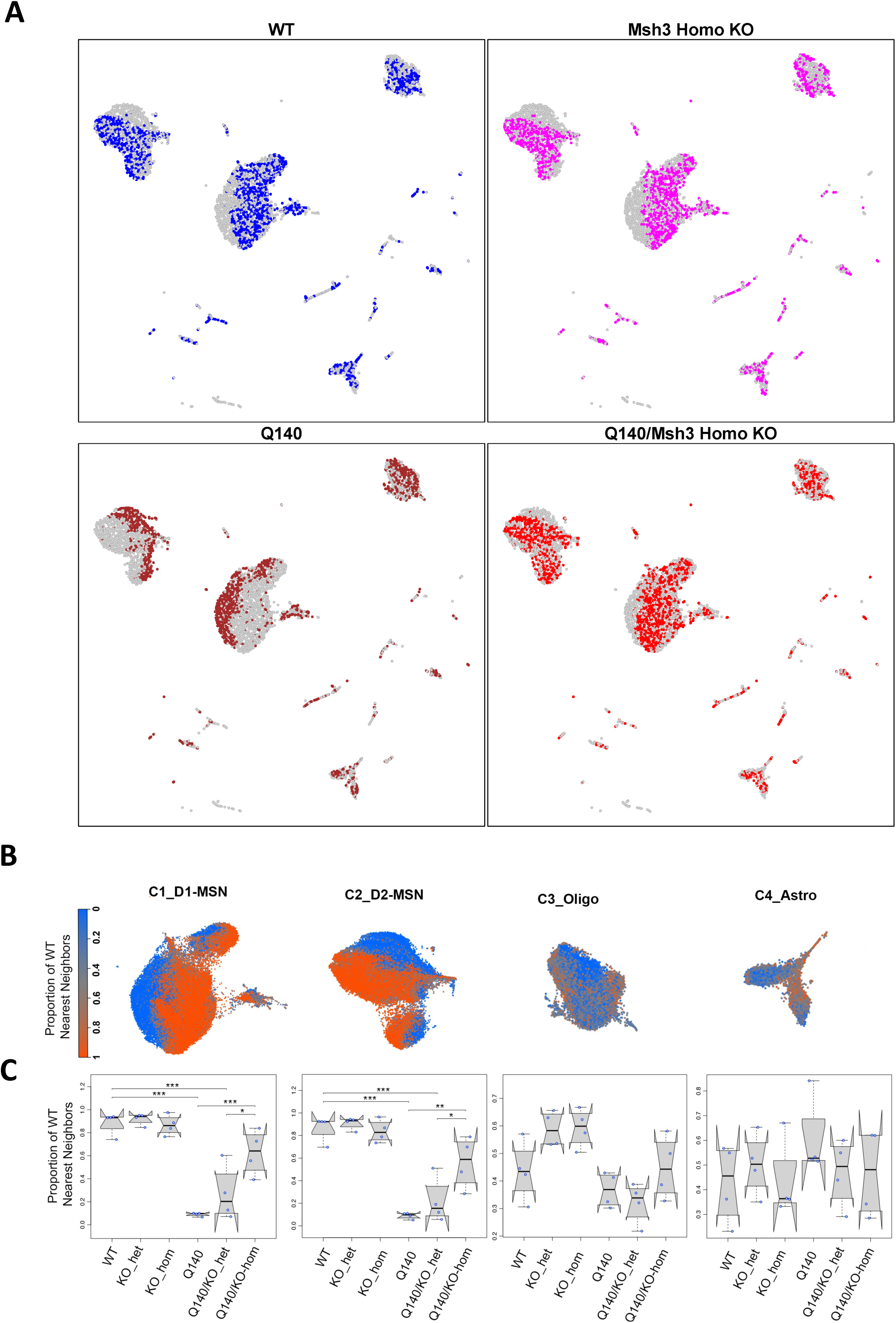
Neighborhood analysis by clusters/genotypes in Msh3 KO snRNA-seq. (A) Representative distribution of cells from each genotype in all clusters. Each panel shows a random sample of cells chosen such that half of the foreground cells are from the genotype indicated in the title of the panel and the rest are from other genotypes. Cells from the selected genotype are colored in one of the 6 colors; cells from other genotypes are colored grey. We chose the 50% frequency to illustrate relative frequency of cells from a single genotype in different parts of the UMAP representation of the clusters. (B) UMAP representations of clusters C1_D1-MSN, C2_D2-MSN, C3_Oligo and C4_Astro colored by the proportion of WT cells among 20 nearest WT or Q140 neighbors. (C) Boxplots of the proportion of WT nearest neighbors across the 6 genotypes in each of the 4 clusters in panel B. For these boxplots, the proportions corresponding to individual cells were aggregated by animal resulting in 4 data points per genotype, shown in blue. Tukey HSD test (n=4 per genotype) was used to assess significance of between-genotype differences in separately in each cluster. *: p<0.05; **: p<0.01; ***: p<0.001

**Supplemental Figure S8.**
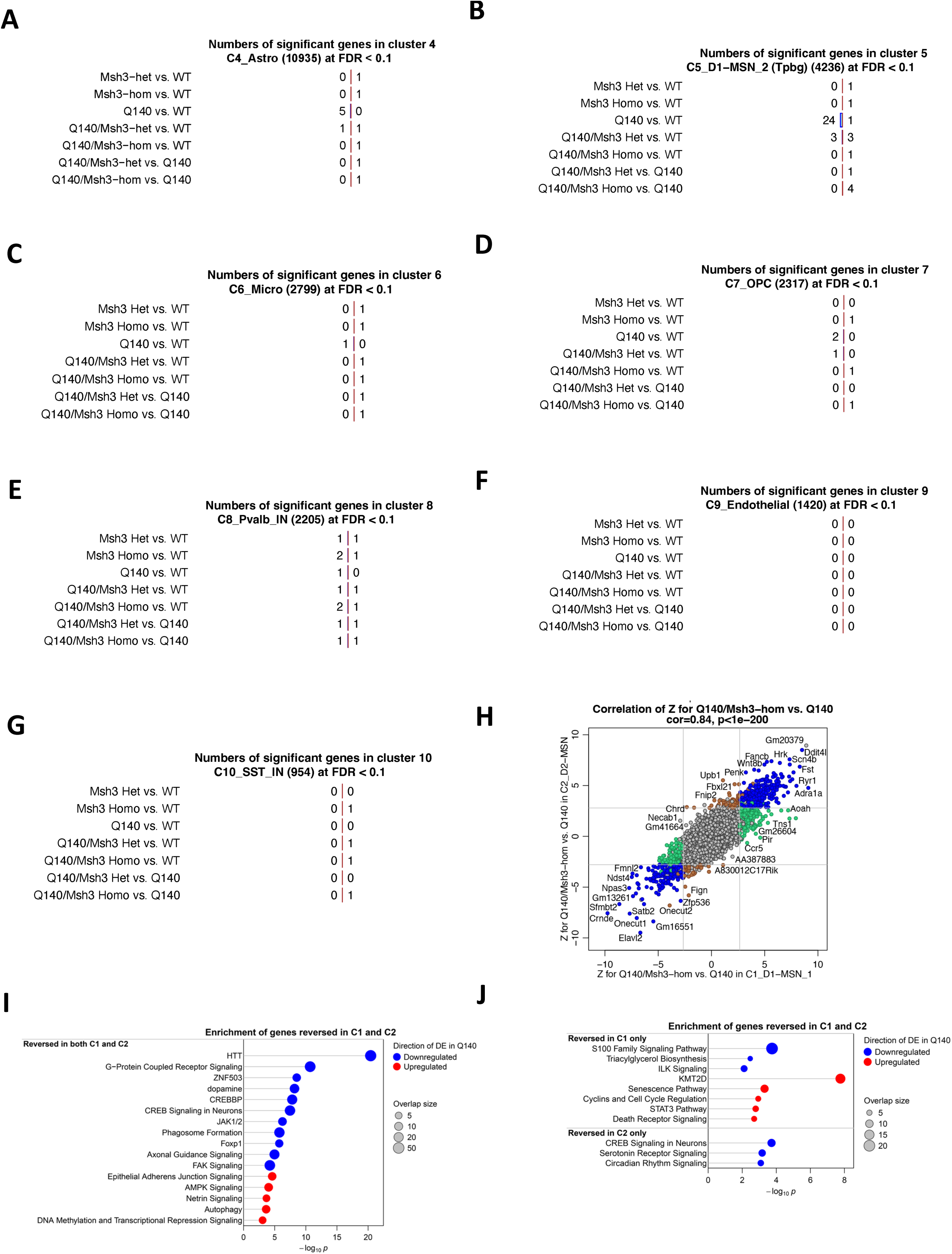
Pseudobulk differential gene expression analysis of Q140/Msh3 KO snRNA-seq. (A) Numbers of DE genes in the astrocyte cluster C4_Astro from pseudobulk analysis of 12m Msh3 KO snRNA-seq. DE genes are identified by pseudobulk analysis at FDR<0.1. The bars are drawn to the same scale as those in Fig. 2D. Blue and red bars represent down- and upregulated genes, respectively. (B) Numbers of DE genes in a sub-population of D1-MSN cluster C5_D1-MSN_2 (Tpbg) from pseudobulk analysis of 12m Msh3 KO snRNA-seq. (C) Numbers of DE genes in the microglia cluster C6_Micro from pseudobulk analysis of 12m Msh3 KO snRNA-seq. (D) Numbers of DE genes in the oligodendrocyte precursor cell cluster C7_OPC from pseudobulk analysis of 12m Msh3 KO snRNA-seq. (E) Numbers of DE genes in the parvalbumin-positive neuron cluster C8_Pvalb_IN from pseudobulk analysis of 12m Msh3 KO snRNA-seq. (F) Numbers of DE genes in the endothelial cell cluster C9_Endothelial from pseudobulk analysis of 12m Msh3 KO snRNA-seq. (G) Numbers of DE genes in the somatostatin-expressing (SST) interneuron cluster C10_SST_IN from pseudobulk analysis of 12m Msh3 KO snRNA-seq. (H) Scatterplot of pseudobulk DE analysis Wald Z for Q140/Msh3-hom vs. Q140 in cluster C2_D2-MSN (y axis) vs. Z statistics for the same test in cluster C1_D1-MSN (x axis). Color indicates genes that are significantly reversed in both clusters (blue), only in C1 (green), only in C2 (brown) and in neither of the clusters (grey). (I) Enrichment p-values for genes commonly reversed in both C1_D1-MSN and C2-D2-MSN in selected terms identified by Ingenuity Pathway Analysis. (J) Enrichment p-values for genes selectively reversed in either C1_D1-MSN or C2-D2-MSN in selected terms identified by Ingenuity Pathway Analysis.

**Supplemental Figure S9.**
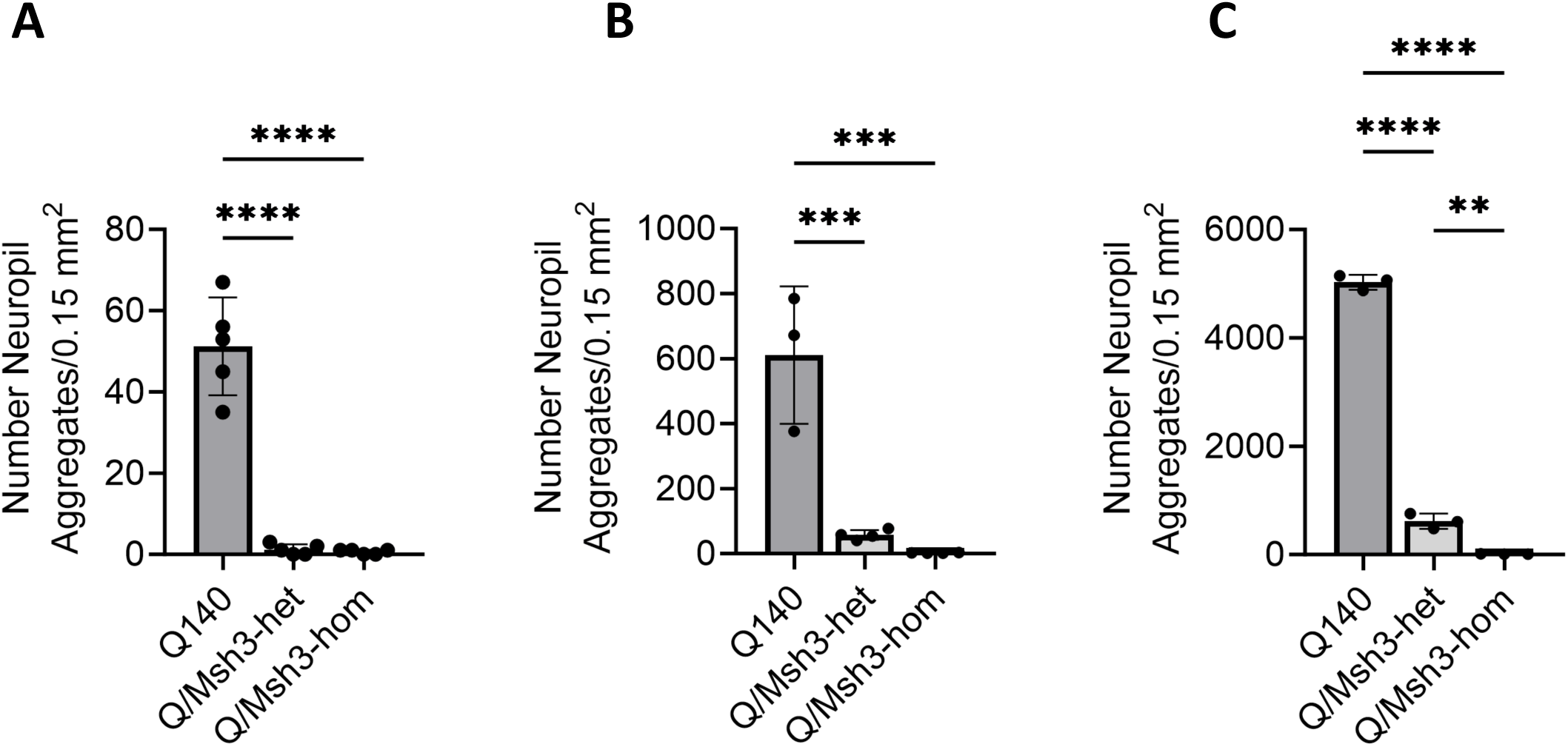
Quantification of mHtt Neuropil aggregate in Q140/Msh3-KO mouse cohort. (A) The numbers of neuropil aggregates in striata of 6m Msh3/Q140 cohort. Data are shown as mean ± SEM, n = 3-6 per genotype. Data analysis by one-way ANOVA with Tukey’s multiple comparison tests: **** p<0.0001, *** p<0.001, ** p<0.01. (B) The numbers of neuropil aggregates in striata of 12m Msh3/Q140 cohort. (C) The numbers of neuropil aggregates in striata of 20m Msh3/Q140 cohort.

**Supplemental Figure S10.**
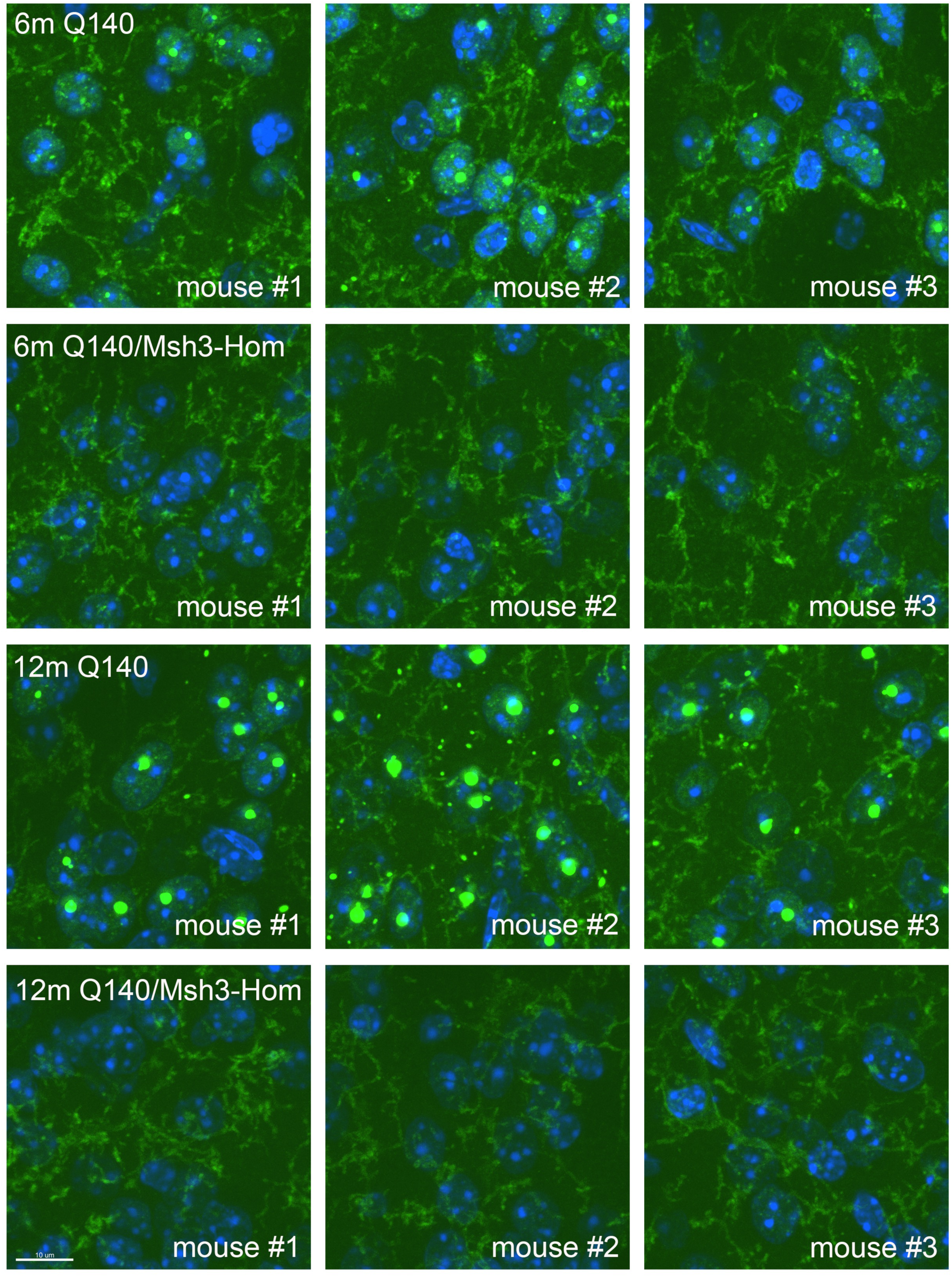
Mutant Huntingtin aggregate staining with EM48 antibody in Q140 and Q140/Msh3-hom mice at 6m and 12m of age. Pathological results were confirmed using Em48 (mHtt, green) and DAPI (nucleus, blue). Three mice of each genotype (Q140 or Q140/Msh3-hom) per age (6m or 12m) were showed. Msh3 KO not only reduces mHtt inclusions, also eliminates diffused nuclear and neuropil mHtt aggregates.

**Supplemental Figure S11.**
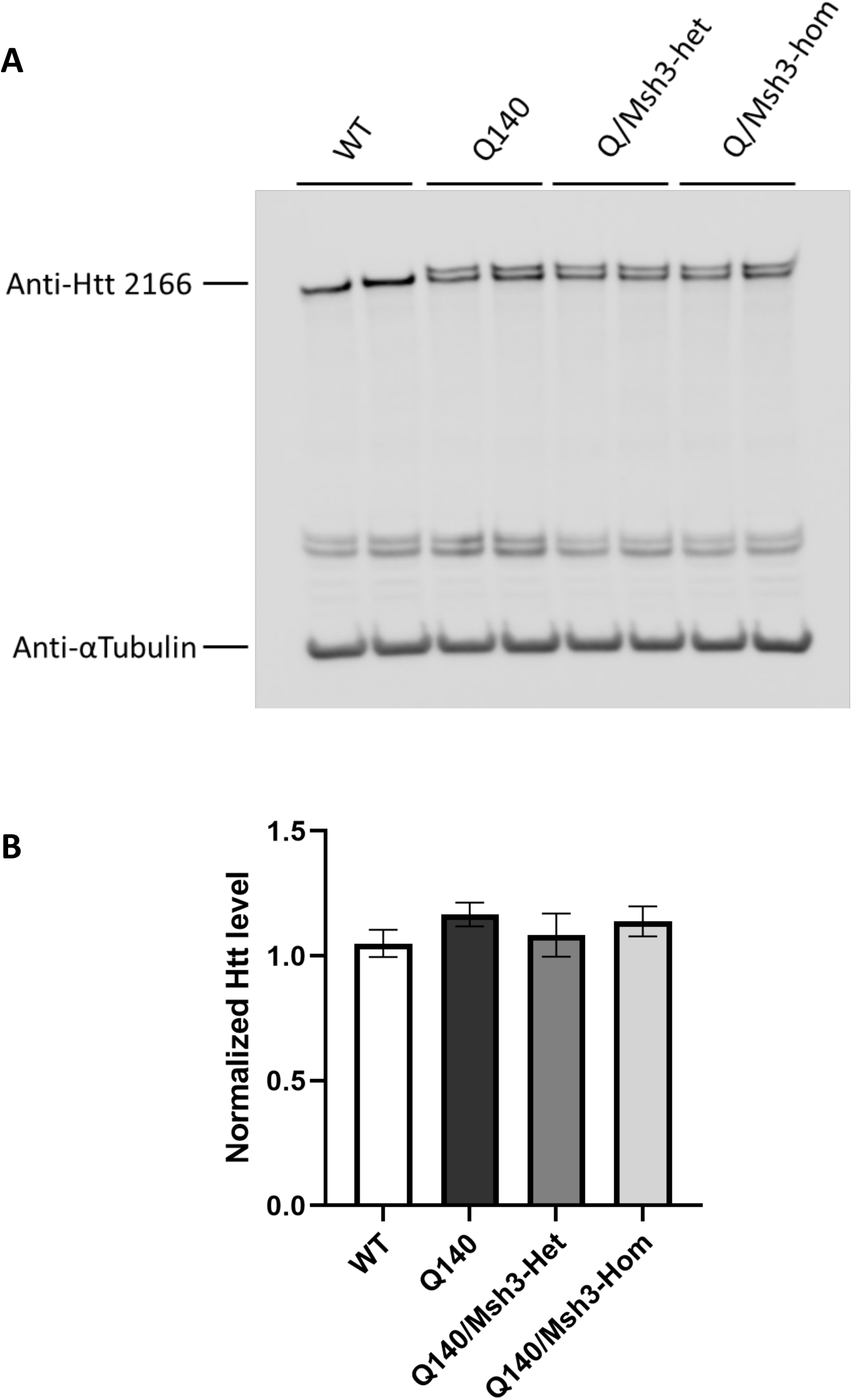
Western blot analysis of wildtype and mutant Htt expression levels in the striata from mice with and without Msh3 KO alleles. (A) Western blots of striatal lysates from 3m WT, Q140, Q140/Msh3-het and Q140/Msh3-hom mice were probed with anti-Htt MAB2166 and anti-αTubulin (loading control). A representative image from two independent experiments is shown. (B) Quantification of total Htt protein levels among the genotypes (from n=4 per genotype).

**Supplemental Figure S12.**
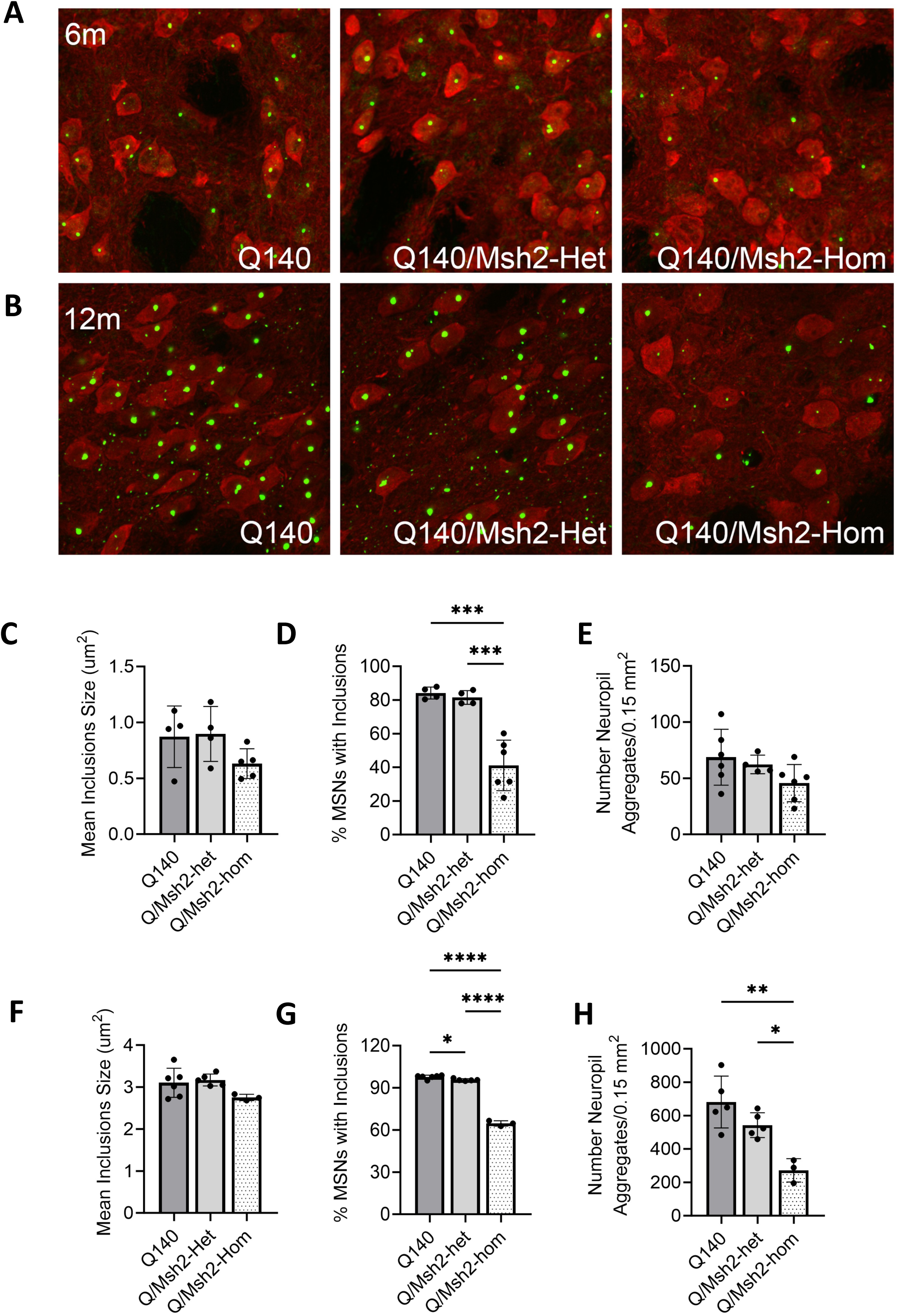
Mutant Htt aggregate pathology in the striata from 6m and 12m Q140/Msh2 KO mouse cohort. (A) Representative images of PHP1-stained mHtt aggregates in striata of 6m Q140/Msh2 KO cohort. Coronal striatal sections (30µm thickness) were double stained with anti-Darpp32 (red) and anti-mHtt (PHP1, green) antibodies. (B) Representative images of PHP1-stained mHtt aggregates in striata of 12m Q140/Msh2 KO cohort. (C) Average size of nuclear inclusion (NI) in striata of 6m Q140/Msh2 KO cohort. For all pathological quantification, data are shown as mean ± SEM, n = 3-6 for each age groups. Data analysis by one-way ANOVA with Tukey’s multiple comparison tests: * p<0.05, ** p<0.01, *** p<0.001, **** p<0.0001. (D) The percentage of medium spiny neurons (MSNs) with NIs in striata of 6m Msh2/Q140 cohort. (E) The numbers of neuropil aggregates in striata of 6m Q140/Msh2 KO cohort. (F) The average size of NIs in striata of 12m Q140/Msh2 KO cohort. (G) The percentage of MSNs with NIs in striata of 12m Q140/Msh2 KO cohort. (H) The numbers of neuropil aggregates in striata of 12m Q140/Msh2 KO cohort.

**Supplemental Figure S13.**
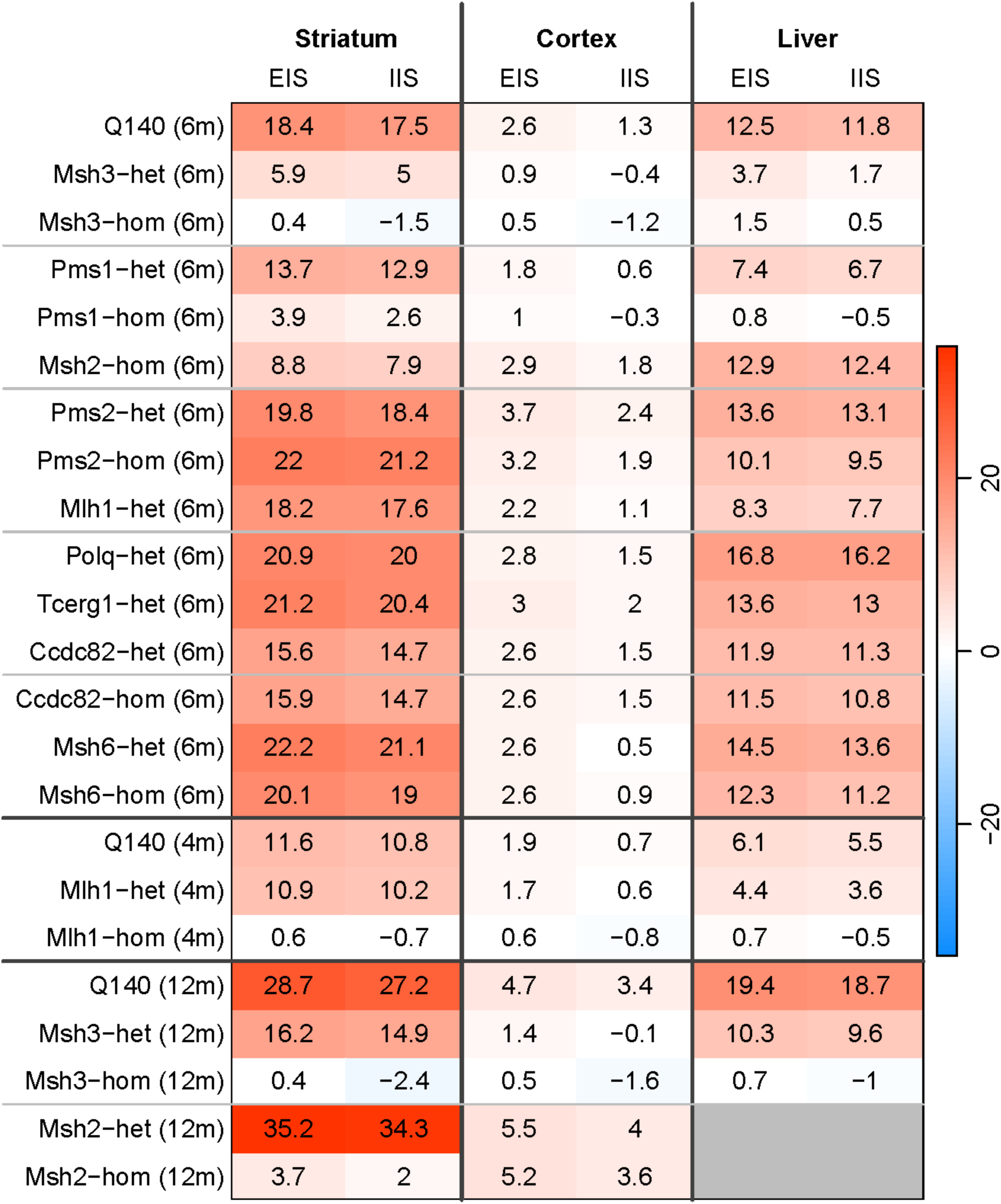
Somatic CAG Instability Indices and Expansion Indices in the striatum, cortex and liver tissues from all KO lines crossed to Q140. The average CAG instability in the striatum, cortex, and liver in all lines. EIS: expansion index score; IIS: instability index score. The values of Q140 are the means of all Q mice at that age from all crossing, all other values are the means of 5-6 mice from Q140 with the indicated perturbation. Red and Blue color indicate instability index increase (positive values) and decrease (negative values), respectively.

**Supplemental Figure S14.**
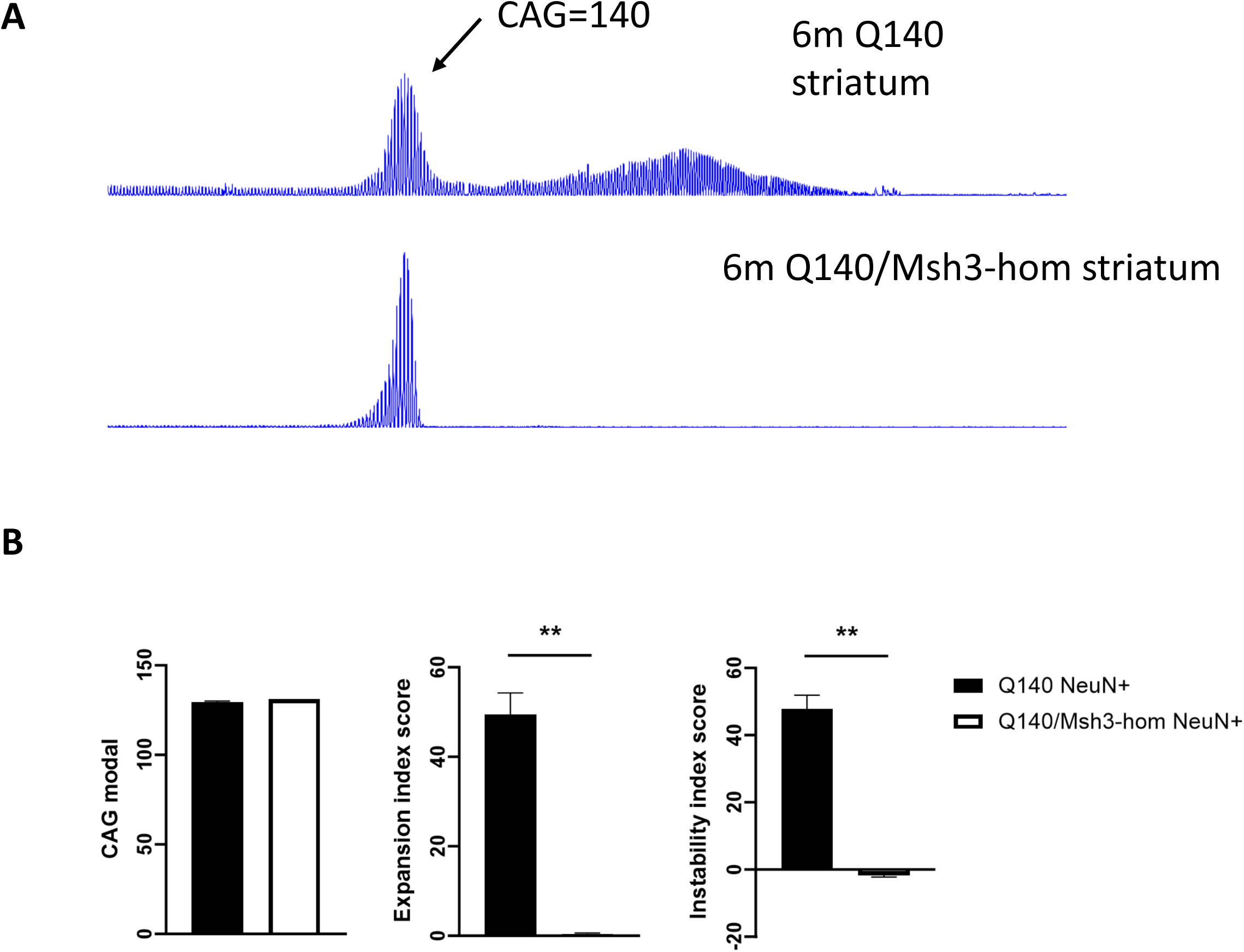
CAG Instability in FANS-purified (anti-NeuN^+^) striatal neuronal nuclei from 6m-old Q140 and Q140/msh3-hom mice. (A) Representative GeneMapper traces of mHtt CAG repeats in neuronal nuclei purified with FANS using antibody against NeuN from 6m Q140 and Q140/Msh3-hom striatal tissues. (B) Somatic CAG instability indices in NeuN+ nuclei of 6m Q140 and Q140/Msh3-hom striatal tissues.

**Supplemental Figure S15.**
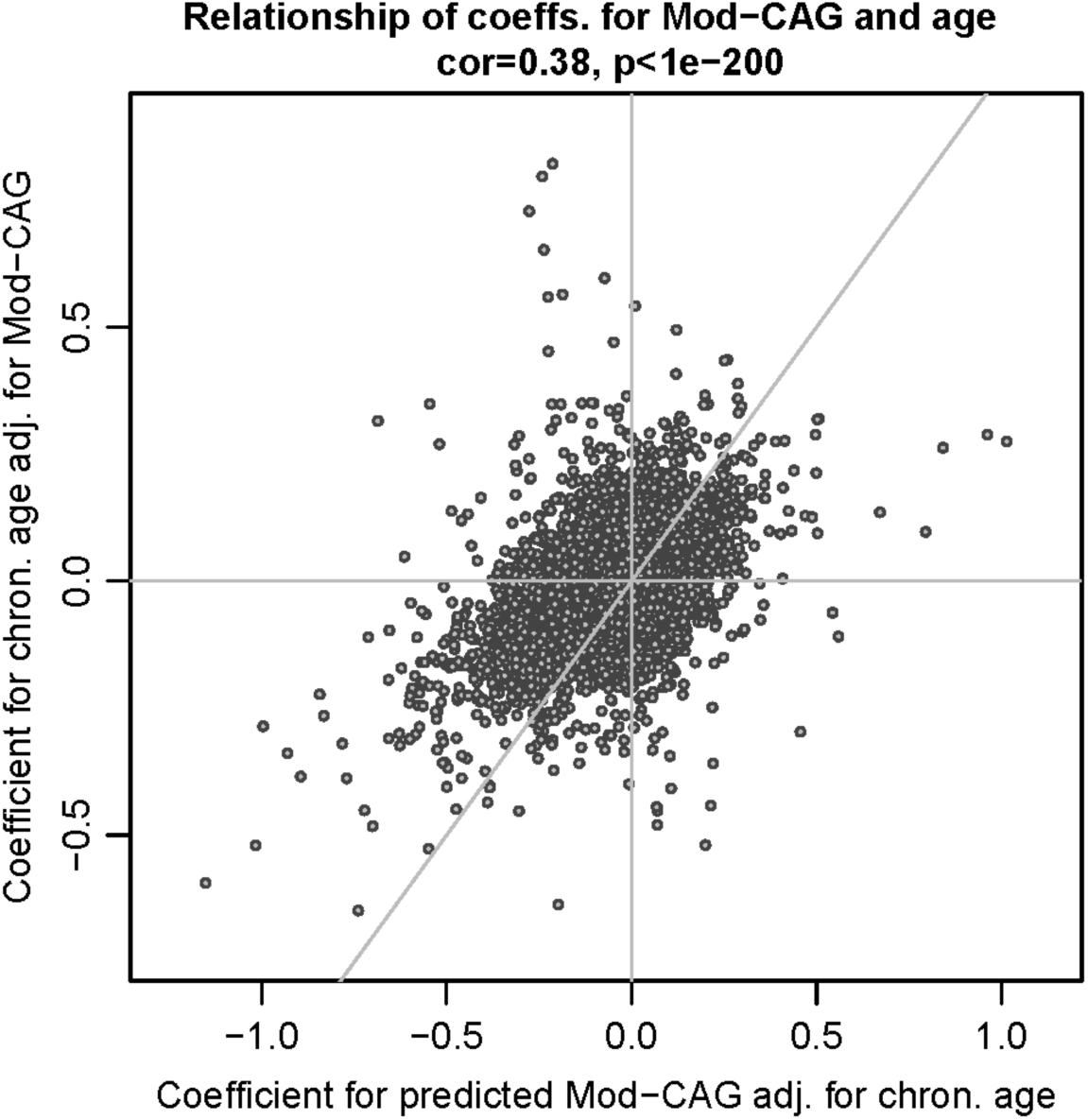
Relationships among model coefficients for Mod-CAG and chronological age. Relationship of gene-wise linear model coefficients for gene expression regressed on Mod-CAG adjusted for chronological age (x-axis) and chronological age adjusted for Mod-CAG (y-axis). The diagonal grey line represents equality x=y. Each point represents a gene.

**Supplemental Figure S16.**
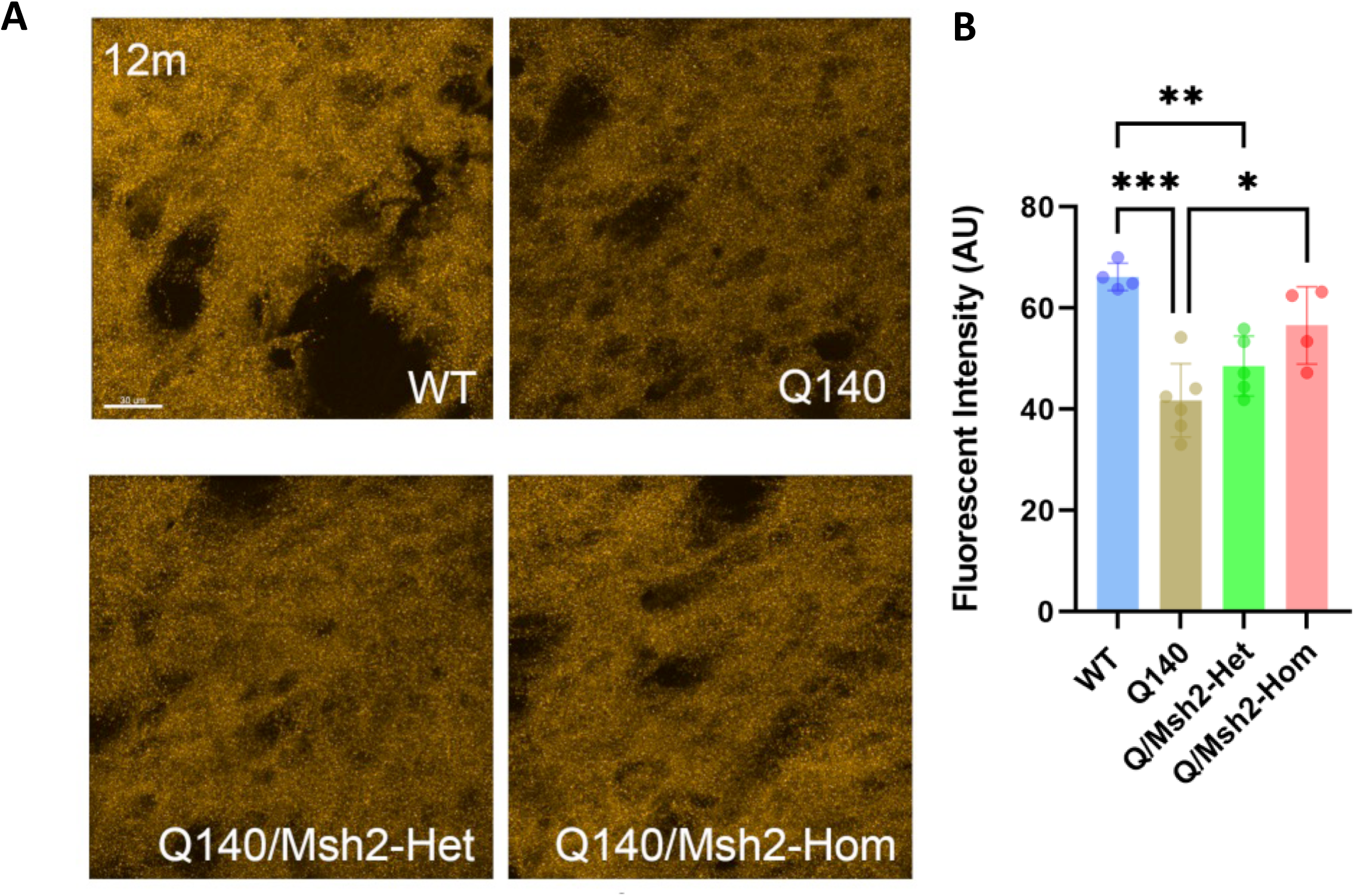
Actn2 level in 12m WT, Q140, Q140/Msh2-het, and Q140/Msh2-hom striata at 12m of age. (A) Representative images of Actn2 (post-synaptic marker) staining in striata of 12m Msh2/Q140 cohorts. Coronal striatal sections (10µm thickness) were stained with anti-Actn2 antibody. (B) Quantifications of Actn2 intensity in striata of 12m Msh2/Q140 cohorts. Results are shown as mean ± SEM from 4 brains for each group. Data analysis by one-way ANOVA with Tukey’s multiple comparison tests: * p<0.05, ** p<0.01, *** p<0.001.

